# Fibrillar adhesions are the primary integrin complexes shaped by matrix topography

**DOI:** 10.1101/2025.11.26.690075

**Authors:** Aaron J. Farrugia, Shao-Zhen Lin, Yee Jun Yuen, Srinivas Sheshagiri Prabhu, Yilin Wang, Mui Hoon Nai, Sebastian Foo, Hamizah A. Cognart, Chwee Teck Lim, Gianluca Grenci, Boon Chuan Low, Pakorn Kanchanawong, Jean-François Rupprecht, Jacques Prost, Alexander D. Bershadsky

## Abstract

Mechanisms of matrix topography recognition are poorly understood. Here, we show that α5β1-integrin mediated fibrillar adhesions serve this function. While on planar substrates, their formation requires fibronectin secretion, tensins, and actomyosin contractility, these requirements are bypassed on nanotopographical features. While focal adhesions avoid these features, fibrillar adhesions rapidly align along pre-existing fibrous cell-derived matrix or electrospun nanofibers where they can then template fibronectin fibrils. Topography-induced fibrillar adhesions depend primarily on α5β1-integrin clustering and disassemble upon cortical stiffening driven by myosin-II overactivation or increased membrane tension. We propose a generic theoretical model where the adhesion receptor favours the membrane and substrate planes to be tilted relative to each other. This model matches experimental observations of preferential α5β1-integrin clustering along nanofibers and concave edges of large negative curvature along micro-ridges. These findings establish fibrillar adhesions as primary adhesion complexes that form independently of focal adhesions in response to matrix topography.

## Introduction

Integrin-mediated adhesion to the extracellular matrix (ECM) regulates cell behaviour during development, tissue homeostasis and disease [1–3]. Upon binding ECM proteins, integrins initiate the formation of diverse multi-protein adhesion complexes including focal adhesions, fibrillar adhesions, podosomes/invadopodia, reticular adhesions, filopodia, curved adhesions and 3D matrix adhesions. Despite often sharing core components such as talin and frequently incorporating F-actin into their plaques, these structures differ markedly in their functions, including their abilities to sense and respond to mechanical cues, remodel the ECM, and mediate cell attachment across 2D and 3D microenvironments [4–11]. The mechanisms driving the formation and specialization of these diverse structures remain insufficiently understood.

Focal adhesions (or FAs) are the best-characterized and widely considered as the archetypal integrin complexes. They arise from nascent focal complexes that mature in a force-dependent, actomyosin-driven manner into micron sized plaques associated with thick F-actin cables (stress fibers) and rapidly disassemble upon inhibition of myosin-II activity [4, 5]. Fibrillar adhesions (or FBs) represent a less understood but functionally important adhesion complexes, particularly in matrix-producing cells such as fibroblasts and endothelial cells. Fibrillar adhesions connect cells to fibronectin (FN) fibrils via α5β1-integrins and are key drivers of fibronectin fibrillogenesis [6, 12–15]. Fibronectin is a crucial ECM component, essential during embryogenesis and strongly associated with developmental and tumour angiogenesis [16, 17]. Fibrillar adhesions are morphologically distinct from focal adhesions, appearing as centrally located, elongated or wavy structures, particularly enriched in tensin proteins (tensin1 and tensin3) and while containing talin, they lack canonical focal adhesion markers such as paxillin and zyxin [6, 12, 13]. While early studies suggested that fibrillar adhesions arise from mature focal adhesions by myosin-II dependent segregation of fibronectin-associated α5β1-integrin from peripheral focal adhesions [12, 13], there are examples of fibrillar adhesions and fibronectin fibrils forming independently of focal adhesions [14]. Unlike focal adhesions, myosin-II activity is dispensable for the maintenance of fibrillar adhesions, since they are apparently insensitive to Rho-Kinase (ROCK) inhibition [12, 15]. Similarly, fibronectin fibrillogenesis has also been proposed to require myosin-II for both secretion [18] and the mechanical unfolding of cryptic binding sites within fibronectin monomers [19–22]. The individual contributions of myosin-II to fibrillar adhesion formation versus fibronectin assembly have not yet been clearly dissected, due to the strong interdependence of these processes.

Here, we show that substrate nano-topography is the major factor determining the assembly of fibrillar adhesions. Unlike focal adhesions which avoid topographical features, fibrillar adhesions rapidly form and align along pre-existing cell-derived matrix fibers and electrospun nanofibers, independently of myosin-II activity and cellular fibronectin deposition. The physical properties of the membrane are crucial as fibrillar adhesion disassemble upon interventions that increase cortical stiffness (such as excessive myosin activation) or membrane tension. Finally, we propose a theoretical model describing a minimal system based on the geometric properties of bent α5β1-integrin, which assumes that the free energy of the adhesion receptor is minimized when the membrane and substrate planes are tilted relative to each other. This model not only accounts for the observed sensitivity to membrane tension but predicts the formation of fibrillar adhesions along nanofibers and concave regions across various microfabricated micro-ridges. These findings position α5β1-integrin and fibrillar adhesions as a focal adhesion-independent principal response to ECM topography.

## Results

### Fibrillar adhesions require fibronectin secretion and myosin-IIA for their assembly on planar glass but not on fibrous cell-derived matrix

Fibrillar adhesions formed by human umbilical vein endothelial cells (HUVECs) appear as centrally located, elongated, straight or wavy structures detected by an antibody targeting α5β1-integrin in extended open conformation (clone SNAKA51) [23, 24], which colocalize with extracellular fibronectin fibrils **(Fig. 1A)**. Remarkably, SNAKA51 positive fibrillar adhesions can also be detected by mAb13 antibody **(Fig. S1A)** interacting with β1-integrin in its bent conformation [25]. These structures are also enriched in other proteins including tensin1, tensin3, talin1, and KANK2 **(Fig. S1A)**. In contrast, focal adhesion localize at F-actin stress fiber termini [26, 27] and contain canonical proteins such as paxillin, vinculin, zyxin and αv-integrin with little to no overlap with fibrillar adhesions **(Fig. S1B).** Tensin1 and tensin3 may occasionally localize to focal adhesions, particularly upon overexpression, however these sites lack associated fibronectin fibrils **(Fig. S1C)**.

**Figure 1.**
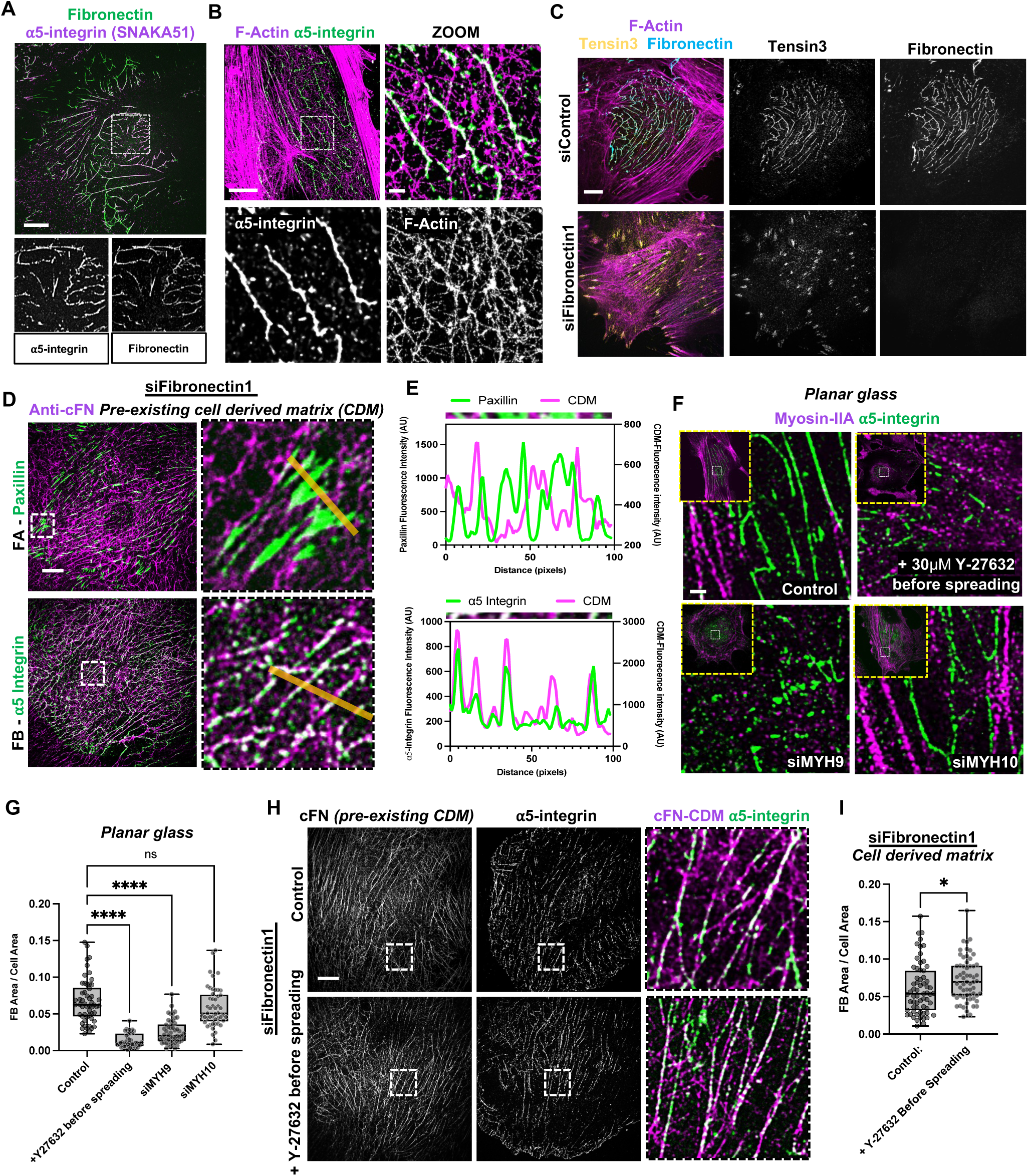
Fibronectin secretion and actomyosin contractility are dispensable for fibrillar adhesions formed on cell derived matrix. (A) Structured illumination microscopy (SIM) image of HUVEC spread overnight on planar glass showing fibronectin (green) and fibrillar adhesions identified by α5-integrin visualized by SNAKA51 antibody recognizing the open, extended conformation (magenta). Insets show single channels of zoomed regions. Scale bar, 10µm. (B) STORM images of fibrillar adhesions (α5-integrin SNAKA51, green) and F-actin (phalloidin, magenta) in HUVEC. Fibrillar adhesions are not associated with thick F-actin cables. Insets highlight fibrillar adhesions alignment with thin F-actin cables and possibly individual filament. Scale bars, 10µm (overview), 1µm (zoom). (C) Control and fibronectin1 depleted (siFN1) HUVEC on planar substrates showing tensin3 (yellow) and external fibronectin fibrils (cyan). Depletion of FN1 results in a loss of fibrillar adhesions and redistribution of tensin3 to focal adhesion sites at stress fiber termini. Corresponding individual channels displayed separately. Scale bar, 10µm. (D) FN1 knockdown cells grown on pre-formed fibrous cell-derived matrix (CDM) stained with an antibody targeting cellular fibronectin, which is specific for cellular and not plasma fibronectin (cFN present in the deposited CDM, magenta). Focal adhesions are visualized with paxillin (FA, green) and fibrillar adhesions with α5-integrin (FB, green). Insets show zoomed regions with yellow lines indicating positions for intensity line scans in E. (E) Line scan plots of fluorescence intensities for paxillin and α5-integrin (green), together with corresponding underlying cellular fibronectin (cFN marking CDM, in magenta) along indicated lines in (D). Focal adhesions avoid fibrous ECM while fibrillar adhesions fully associate with it. (F) Cells plated on planar glass under control conditions or in the presence of 30µM ROCK inhibitor Y-27632 and RNAi depletion of myosin-IIA (siMYH9) or myosin-IIB (siMYH10), showing myosin-IIA heavy chain (magenta) and α5-integrin (green). Insets show the cell images with low magnification. Formation of fibrillar adhesions is suppressed by treatment with Y-27632 or myosin-IIA knockdown. Scale bar, 1µm. (G) Quantification of the ratio of fibrillar adhesion area to total cell area (FB Area/Cell Area) under conditions specified in (F), siControl (*n*=54), +Y-27632 before spreading (*n*=45), siMHY9 (*n*=51), siMHY10 (*n*=50), *N*=3, box-and-whisker plots, whiskers extend from the minimum to maximum values, the box extends from the 25th to the 75th percentile and the line within the box represents the median, *P*-values calculated using one way ANOVA, with each mean compared to the mean of the control (****, *P* < 0.0001, ns = not significant). (H) HUVEC under fibronectin knockdown (siFN1) grown on pre-existing CDM with or without treatment with 30µM Y-27632 before cell spreading, showing cFN in the pre-existing CDM and α5-integrin in fibrillar adhesions. Insets are shown with high magnification in the right column display merged regions (cFN – magenta, α5-integrin – green). Note that fibrillar adhesions are assembled along CDM fibers even in a ROCK independent fashion. Scale bar, 10µm. (I) Quantification of ratio of fibrillar adhesion area to total cell area (FB Area/Cell Area) for siFN1 HUVEC without treatment (*n*=63) or treated with 30µM Y-27632 before spreading (*n*=61), *N*=3, box-and-whiskers plot (min, median, max) with all data points shown, *P*-values calculated using two-tailed unpaired t-test (*, *P* < 0.05).

To examine the relationship between fibrillar adhesions and the actin cytoskeleton we used stochastic optical reconstruction microscopy (STORM) [28]. This revealed that fibrillar adhesions associate with thin F-actin cables or possibly individual filaments **(Fig. 1B)**, but do not overlap with thick stress fibers. A known distinction between focal adhesions and fibrillar adhesions is their differential sensitivity to inhibitors of myosin-II activity: pre-formed fibrillar adhesions are resistant to Rho-kinase (ROCK) inhibition by Y-27632, which blocks myosin-II filament assembly, while focal adhesions rapidly disassemble [12]. We confirmed this by live-cell imaging of GFP-tensin3 labeled fibrillar adhesions and mCherry-paxillin labeled focal adhesions during ROCK inhibition **(Supplemental Movie 1)**. Fibrillar adhesion formation on planar glass has been reported to fully depend on cellular fibronectin secretion [13, 29]. Consistently, we observed that knockdown of *FN1* by RNAi (siFN1) **(Fig. S1D)** eliminates fibrillar adhesions and causes tensin3 to re-localize to focal adhesion sites at the termini of stress fibers **(Fig. 1C)**.

To examine fibrillar adhesions in a more physiologically relevant context, we generated a cell-derived matrix (CDM) by culturing HUVECs on gelatin-coated coverslips followed by decellularization using a detergent to retain the secreted ECM [30] **(Fig. S1E)**. We then plated fresh HUVEC on top of the pre-formed CDM and observed that focal adhesion-associated proteins such as paxillin **(Fig. 1E)** and vinculin **(Fig. S1G, H)** preferentially localized to regions largely devoid of fibrous ECM **(Fig. S1F)**. When fibronectin secretion was blocked by FN1 knockdown, fibrillar adhesions still formed and aligned precisely along pre-existing fibrous cellular fibronectin in the CDM **(Fig. 1D, E)**. This was not due to increased ligand availability as coating planar glass substrates with excess human plasma fibronectin failed to induce fibrillar adhesion formation **(Fig. S1I)**.

It has been previously shown that fibrillar adhesion formation on planar substrates depends on myosin-II activity as it can be suppressed by blebbistatin treatment [15]. We confirmed that myosin activity is required as ROCK inhibition prior to cell spreading on planar substrates impaired fibrillar adhesion elongation and reduced their abundance **(Fig. 1F, G)** as well as fibronectin fibrillogenesis **(Fig. S1J).** RNAi knockdown experiments further showed that fibrillar adhesion formation is specifically dependent on myosin-IIA (siMYH9), but not myosin-IIB (siMYH10) **(Fig. 1F, G, S1K)**. We further evaluated the role of myosin activity and pre-existing focal adhesions upon cell spreading on CDM. HUVECs depleted of FN1 were plated onto CDMs in the presence of ROCK inhibitor. Despite inhibited contractility and the absence of mature focal adhesions, cells readily formed numerous fibrillar adhesions that closely recapitulated the architecture of the underlying ECM **(Fig. 1H, I)**. These findings demonstrate that cellular fibronectin secretion, ROCK dependent myosin-IIA driven contractility, and the presence of mature focal adhesions are dispensable for fibrillar adhesion formation when cells are provided with a pre-assembled fibrous cell-derived matrix.

### Fibrillar adhesions, but not focal adhesions, associate with electrospun nanofibers

To clarify the role of fibrillar nano-topography in the formation of fibrillar adhesions, we prepared substrates consisting of electrospun nanofibers made from thermoplastic polyurethane (TPU) [31], mimicking the nanoscale ECM structure. These fibers were spun onto glass coverslips, forming a randomly oriented meshwork with an average fiber diameter of approximately 200 nm **(Fig. 2A)**, resembling the dimensions of native ECM fibrils. The substrates were then coated with human plasma fibronectin to facilitate cell adhesion and to provide relevant integrin ligands. When HUVECs were cultured on these nanofiber substrates we observed a striking spatial segregation of adhesion complexes. Classical focal adhesion proteins such as paxillin **(Fig. 2B, C)**, vinculin, zyxin, and αv-integrin **(Fig. S2A, C)** were predominantly localized to the planar, flat regions between the nanofibers, with minimal association along the fibers themselves. This distribution is consistent with our observations in fibrillar CDMs, where focal adhesions preferentially form in areas devoid of ECM fibrils **(Fig. 1D, E, S1F)**. Meanwhile all fibrillar adhesion-associated proteins: extended α5-integrin **(Fig. 2B, C)**, bent β1-integrin, tensin1, tensin3, talin1, and KANK2 **(Fig. 2C, S2A, C)** displayed a perfect alignment along the nanofibers.

**Figure 2.**
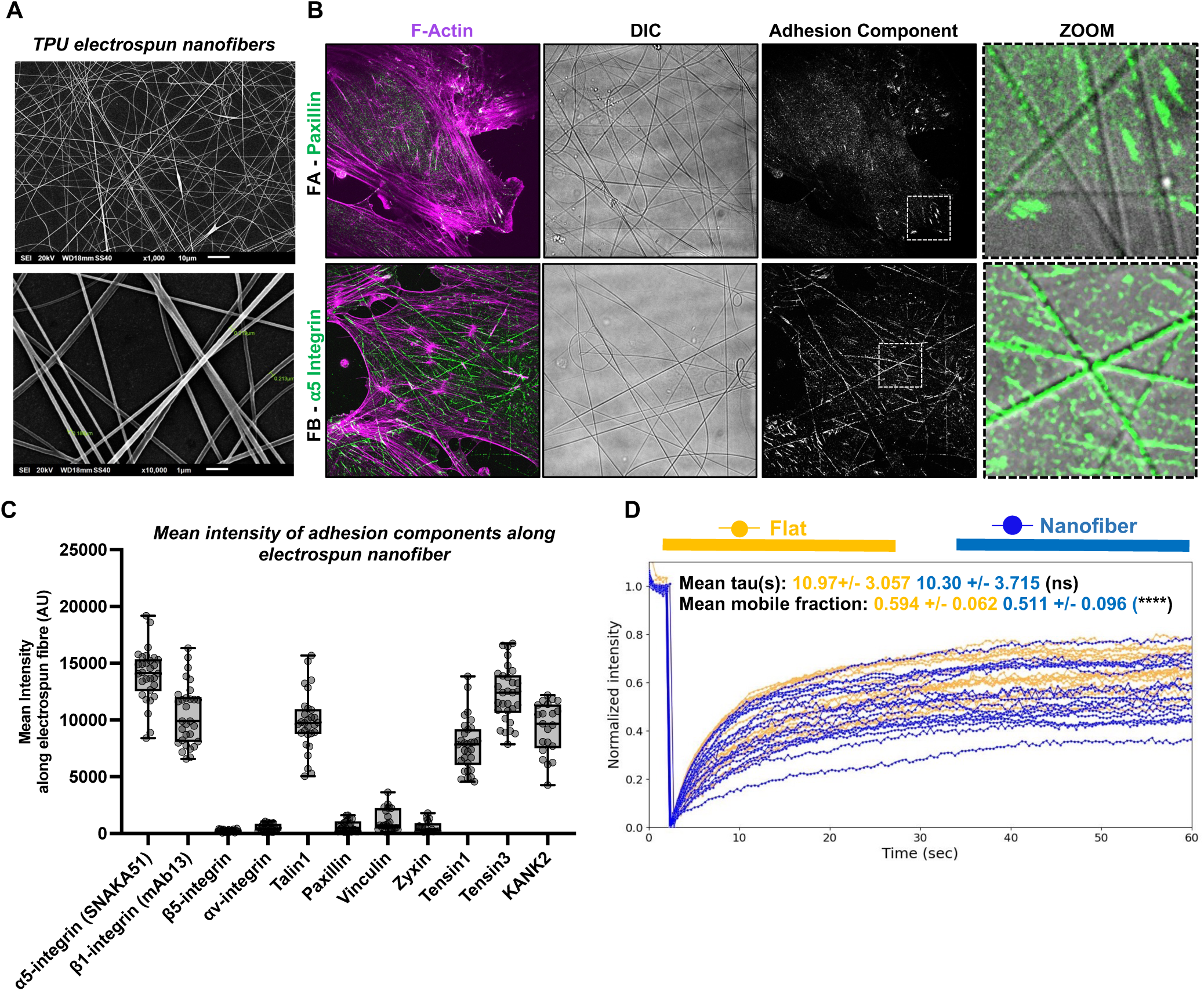
Fibrillar adhesions, but not focal adhesions, assemble along electrospun nanofibers. (A) Scanning electron micrographs of thermoplastic polyurethane (TPU) nanofibers on top of glass coverslips made by electrospinning. Scale bars indicated on images. (B) HUVEC plated on human plasma fibronectin–coated electrospun nanofibers showing F-actin (magenta), focal adhesions visualized with paxillin (FA, green, seen as white at stress fiber termini) and fibrillar adhesions visualized with α5-integrin (FB, green). DIC channel shows nanofibers spun on planar glass. Insets highlight fibrillar adhesion formation along the nanofibers while focal adhesions form on adjacent planar regions. Scale bar, 10µm. (C) Semiquantitative estimation of mean intensities of adhesion components along electrospun nanofibers: α5-integrin SNAKA51 (*n*=30), β1-integrin mAb13 (*n*=30), β5-integrin (*n*=20), αv-integrin (*n*=25), talin1 (*n*=21), paxillin (*n*=22), vinculin (*n*=23), zyxin (*n*=19), tensin1 (*n*=30), tensin3 (*n*=30) and KANK2 (*n*=21). Box-and-whiskers plot (min, median, max) with all data points shown. (D) Quantification of fluorescence intensity recovery after photobleaching of GFP-tensin3 positive fibrillar adhesions on flat (planar) regions (orange) or along electrospun nanofiber regions (blue). Each trajectory represents the average normalized intensity change for adhesions in one cell, with mean exponential time constant, τ (s) and mobile fractions shown on graph (*n*=18 cells with 2-5 individual adhesions examined for each region, *N*=3, Wilcoxon signed-rank test, **** *P* < 0.0001, ns = not significant).

Given this topography-dependent behavior, we hypothesized that curvature-sensing mechanisms might underlie fibrillar adhesion formation. BAR domain-containing proteins are well-known sensors of membrane curvature, with specific subclasses (F-BAR, I-BAR) detecting convex or concave membrane deformations. A previous study has implicated the F-BAR protein FCHo2 in αvβ5-integrin-positive adhesions of cells to nanopillars coated with vitronectin [10]. However, we observed no specific localization of fluorescently tagged FCHo2, or other BAR domain-containing proteins we tested (F-BAR: FCHo1, CIP4, and I-BAR: IRSp53 **(Fig. S2B)**, nor β5-integrin **(Fig. 2C, S2A)**, along electrospun nanofibers.

To gain insight into the dynamic behavior of fibrillar adhesions on nanofibers versus flat regions, we performed fluorescence recovery after photobleaching (FRAP) experiments using GFP-tensin3 as a marker of fibrillar adhesions. We measured the recovery kinetics of GFP-tensin3 fluorescence within fibrillar adhesions located on nanofibers compared to those on planar glass. The recovery time constants (τ) were similar across both conditions **(Fig. 2D, S3A, B, Supplemental Movie 2)**, while mobile fraction was slightly lower on nanofibers **(Fig.S3C.)** Thus, the turnover of fibrillar adhesion components on fibers and planar substrates was comparable.

### Fibrillar adhesion formation along nanofibers, but not fibronectin fibrillogenesis, is independent of actomyosin contractility

To investigate the kinetics of fibrillar adhesion association with nanofibers and its dependence on cellular fibronectin secretion, we examined early cell spreading events. On planar glass substrates, at 30 minutes post-plating, fibrillar adhesions and fibronectin deposition were minimal or absent **(Fig. 3A)**. In stark contrast, fibrillar adhesions rapidly formed along nanofibers within the first 30 minutes of cell attachment, preceding any detectable fibronectin secretion or fibrillogenesis **(Fig. 3A)**. After overnight spreading, cells deposited cellular fibronectin extensively both along nanofibers and across planar regions between fibers **(Fig. 3B)**. When cellular fibronectin secretion was suppressed via siRNA-mediated knockdown of FN1, fibrillar adhesions still formed along nanofibers but were abolished on planar areas in between **(Fig. 3B, C)**. These results demonstrate that fibrillar adhesion formation on nanofibers is rapid and can occur independently of endogenous fibronectin secretion.

**Figure 3.**
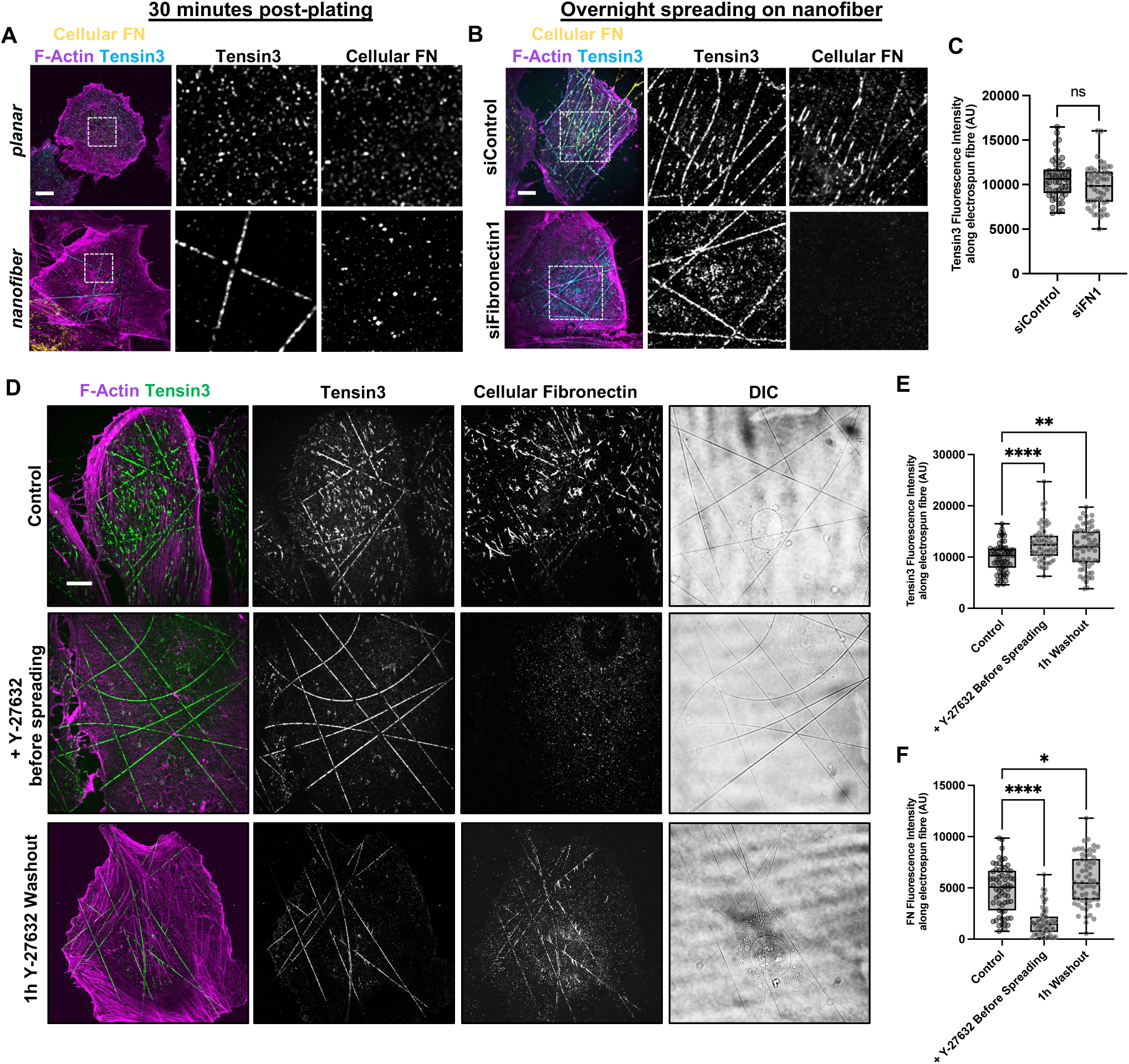
Topography induced fibrillar adhesions form in an actomyosin independent manner and can template myosin-II dependent fibronectin fibrillogenesis. (A) HUVEC fixed 30 minutes after plating on either planar glass or electrospun nanofibers, showing F-actin (magenta), tensin3 (cyan), and cellular fibronectin (yellow). Tensin3 rapidly clusters along nanofibers (but not on planar glass) prior to endogenous fibronectin secretion. Greyscale images show the boxed areas with higher magnification. Scale bar, 10µm. (B) Images of control and FN1 knockdown cells plated overnight on electrospun nanofibers, showing for F-actin (magenta), tensin3 (cyan) and cellular fibronectin (yellow). Tensin3 clusters along electrospun nanofibers (but not adjacent planar regions) even in the absence of fibronectin secretion. Greyscale images show the boxed areas with higher magnification. Scale bar, 10µm. (C) Quantification of mean intensity of tensin3 along nanofibers in control (*n*=49) or fibronectin knockdown cells (siFN1) (*n*=49), *N*=3, box-and-whiskers (min, median, max) graph with all data points shown, *P*-values calculated using two-tailed unpaired t-test, ns = not significant). (D) Images of cells plated overnight on electrospun nanofibers under control conditions (upper row), in the presence of 30µM Y-27632 (middle row), and after washing out the inhibitor for 1 hour. Cells were stained for F-actin (magenta), tensin3 (green), and cellular fibronectin (grey). Tensin3 association with nanofibers persists even under treatment with Y-27632 (although not on adjacent planar regions), while fibronectin deposition is largely inhibited. Cellular fibronectin fibrils reappear along pre-existing tensin3 adhesions on nanofibers after washout of Y-27632. DIC shows nanofibers. Scale bar, 10µm. (E and F) Quantification of mean intensities of tensin3 (E) or cellular fibronectin (F) along nanofibers in control, in the presence of 30µM Y-27632, or after washout of Y-27632 for 1 hour (*n*=62 for each condition, *N*=3, box-and-whiskers plot (min, median, max) with all data points shown, one way ANOVA with each mean compared to the mean of the control, * P < 0.05, ** P < 0.01, **** P < 0.0001).

In agreement with our observations with cells grown on CDMs, the assembly of fibrillar adhesions on nanofibers was also fully independent of myosin activity, when focal adhesion maturation was completely blocked. Addition of ROCK inhibitor before spreading did not inhibit fibrillar adhesion formation on nanofibers **(Fig. 3D, E, S3D)** and, in fact, appeared to enhance them **(Fig. 3E)**. Meanwhile no adhesions were able to form on planar regions between the nanofibers **(Fig. 3D)**. Unlike fibrillar adhesion formation along nanofibers, the process of fibrillogenesis however is ROCK/myosin-II dependent as addition of Y-27632 before spreading strongly inhibited any deposition of fibronectin **(Fig.3D, F)** The treatment allowed us to uncouple nanofiber-guided fibrillar adhesion assembly from fibronectin deposition/fibrillogenesis. After washing out the inhibitor we observed a preferential reassembly of fibronectin fibrils along preformed fibrillar adhesions **(Fig. 3D, F)**, beginning at 30 minutes after washout and becoming very prominent by 60 minutes **(Fig. S3E)**. This data suggests that fibrillar adhesions formed on nanofibers are functional and capable to serve as templates for fibronectin fiber assembly. To test whether this independence of fibrillar adhesion formation on actomyosin contractility extends to other cell types beyond HUVECs, we examined another fibrillar adhesion forming cell type, human foreskin fibroblasts (HFFs). HFFs formed fibrillar adhesions and fibronectin fibrils on planar substrates that were sensitive to ROCK inhibition prior to spreading **(Fig.S3F)**. At the same time, fibrillar adhesions assembled along nanofibers even in the presence of ROCK inhibitor **(Fig.S3G)**, confirming that topography-guided fibrillar adhesion formation is an actomyosin contractility independent process also in fibroblast type cells.

### Clustering of α5β1-integrin along nanofibers is tensin independent

We next focused on the intracellular machinery involved in fibrillar adhesion assembly. Components of fibrillar adhesions, tensin3, tensin1 and KANK1 can undergo liquid–liquid phase separation (LLPS) [32–35]. Treatment of HUVEC on nanofibers with 1,6-Hexanediol, which disrupts phase-separated condensates by interfering with weak hydrophobic interactions [36] led to quantitative extraction of tensin1, suggesting that tensin in fibrillar adhesions is indeed phase separated. At the same time, the tensin1 extraction was not accompanied by removal of α5-integrin assembled along nanofibers **(Fig. 4A, B)**, suggesting that integrin clustering may not depend on LLPS-mediated tensin assembly. To directly test the requirement for tensins, we depleted tensin1 and tensin3 (TNS1 & TNS3) via RNA interference **(Fig. 4C)** and compared formation of fibrillar adhesions (α5-integrin positive clusters) by tensin depleted cells on planar substrates and nanofibers. On plasma fibronectin-coated planar coverslips, TNS1&3 knockdown resulted in a loss of fibrillar adhesions **(Fig. S4A)**. However, when cells were plated on nanofibers, HUVEC depleted of TNS1&3 still formed α5-integrin clusters **(Fig. 4D, E)**, confirming that clustering α5-integrin along topographical features does not require tensin. In contrast, while knockdown of α5-integrin (siITGA5) still permitted cell spreading **(Fig. S4A)**, it inhibited any clustering of tensin3 along nanofibers **(Fig. 4D, E).**

**Figure 4.**
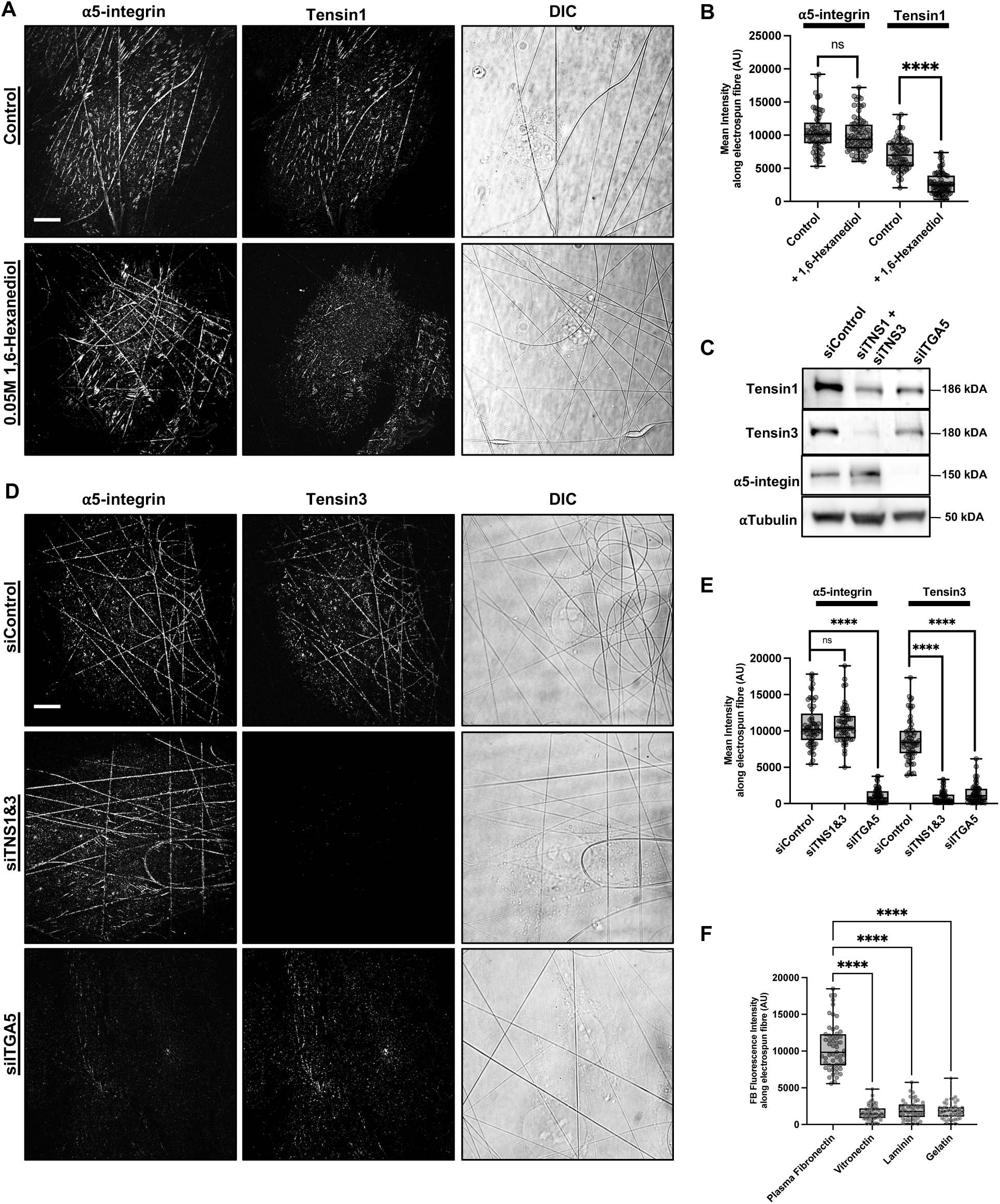
Clustering of α5β1-integrin along nanofibers is tensin independent. (A) Images of HUVEC plated overnight on electrospun nanofibers (upper row) and similar cell after extraction with 1,6-Hexanediol (0.05 M, 5 min). α5-Integrin (SNAKA51) and tensin1 visualized, and DIC showing nanofibers. Scale bar, 10 µm. (B) Quantification of mean intensity of α5-integrin and tensin1 in fibrillar adhesions formed along electrospun nanofibers under control conditions *(n*=69) and after 5 min extraction with 0.05 M 1,6-Hexanediol (*n*=73). Amount of tensin1 is significantly reduced while α5-integrin along nanofibers is preserved (*N*=3, box-and-whiskers plot (min, median, max) with all data points shown, two-tailed unpaired t-test comparing means of α5-integrin or tensin1 average intensity in control or 1,6-Hexanediol treated cells, ns = not significant, **** *P* < 0.0001). (C) Western blot showing protein expression of tensin1 (TNS1), tensin3 (TNS3), and α5-integrin (ITGA5) in siControl, siTNS1&3, and siITGA5 cells. Alpha-tubulin serves as a loading control. Respective molecular weights indicated. (D) Control, tensin1 and 3 double knockdown (siTNS1&3), and α5-integrin knockdown (siITGA5) cells plated overnight on electrospun nanofibers. Images show α5-integrin and tensin3, with DIC to visualize nanofibers. Integrin-α5 localization along nanofibers is not perturbed after depletion of tensin1 and 3, but no localization of tensin3 is observed after depletion of α5-integrin. Scale bar, 10 µm. (E) Quantification of mean intensity of α5-integrin and tensin3 on electrospun nanofibers in RNAi control (*n*=57), siTNS1&3 *(n*=56), and siITGA5 (*n*=56) HUVEC. Box-and-whiskers plot (min, median, max) with all data points shown, one way ANOVA comparing means of α5-integrin or tensin1 mean intensity in siTNS1&3 or siITGA5 cells with those of control, ns = not significant, **** *P* < 0.0001. (F) Quantification of mean intensity of tensin3 clustered along electrospun nanofibers in cells fixed 30 minutes after plating. The electrospun nanofibers were coated with either 10 µg/mL human plasma fibronectin (*n*=55) or 20 µg/mL vitronectin (*n*=51) or 10 µg/mL laminin (*n*=46) or 0.1% gelatin (*n*=42). Only human plasma ficu1ebronectin coating promotes localization of tensin3 along nanofibers. *N*=3, box-and-whiskers plot (min, median, max) with all data points shown, one way ANOVA, **** *P* < 0.0001.

The essential role of α5-integrin in fibrillar adhesion formation was further confirmed using ligand-selective coatings. Integrin-α5β1 exhibits a high degree of specificity for the RGD (Arg-Gly-Asp) motif and the synergy site of fibronectin [37, 38]. To determine whether other extracellular matrix ligands and their corresponding integrins could also support fibrillar adhesion assembly on nanofibers, fibers were also coated with vitronectin, laminin, or gelatin, each engaging multiple integrin type but not α5β1 **(Fig. S4B)**. All coatings supported cell attachment **(Fig. S4C)**, however only cells plated on plasma fibronectin-coated nanofibers formed tensin3-positive fibrillar adhesions within 30 minutes after plating, whereas this was not observed on fibers coated with other ligands **(Fig. 4F, S4C)**. This indicates that α5β1-integrin engagement with fibronectin is specifically required for topography-guided fibrillar adhesion formation.

### Fibrillar Adhesions are sensitive to cortical stiffening driven by overactivation of myosin-II and increased membrane tension

Since fibrillar adhesions appear to be tightly associated with substrate topography, we hypothesized that they may result from externally induced membrane deformation. If this is the case, then conditions that interfere with membrane deformability should disrupt fibrillar adhesion integrity. To test this, we examined how various perturbations known to alter membrane or cortical tension affect fibrillar adhesion stability. First, we subjected cells to mechanical stretching. HUVECs were plated on plasma fibronectin coated elastic polydimethylsiloxane (PDMS) substrates, as previously described [39]. Cells were then exposed to a rapid 16% uniaxial stretch and incubated for 10 minutes, after which they were fixed and stained for fibrillar adhesions (α5-integrin) and focal adhesions (vinculin) **(Fig. 5A)**. Compared to unstretched controls, stretched cells exhibited significant disassembly of fibrillar adhesions, while focal adhesions remained intact **(Fig. 5B)**.

**Figure 5.**
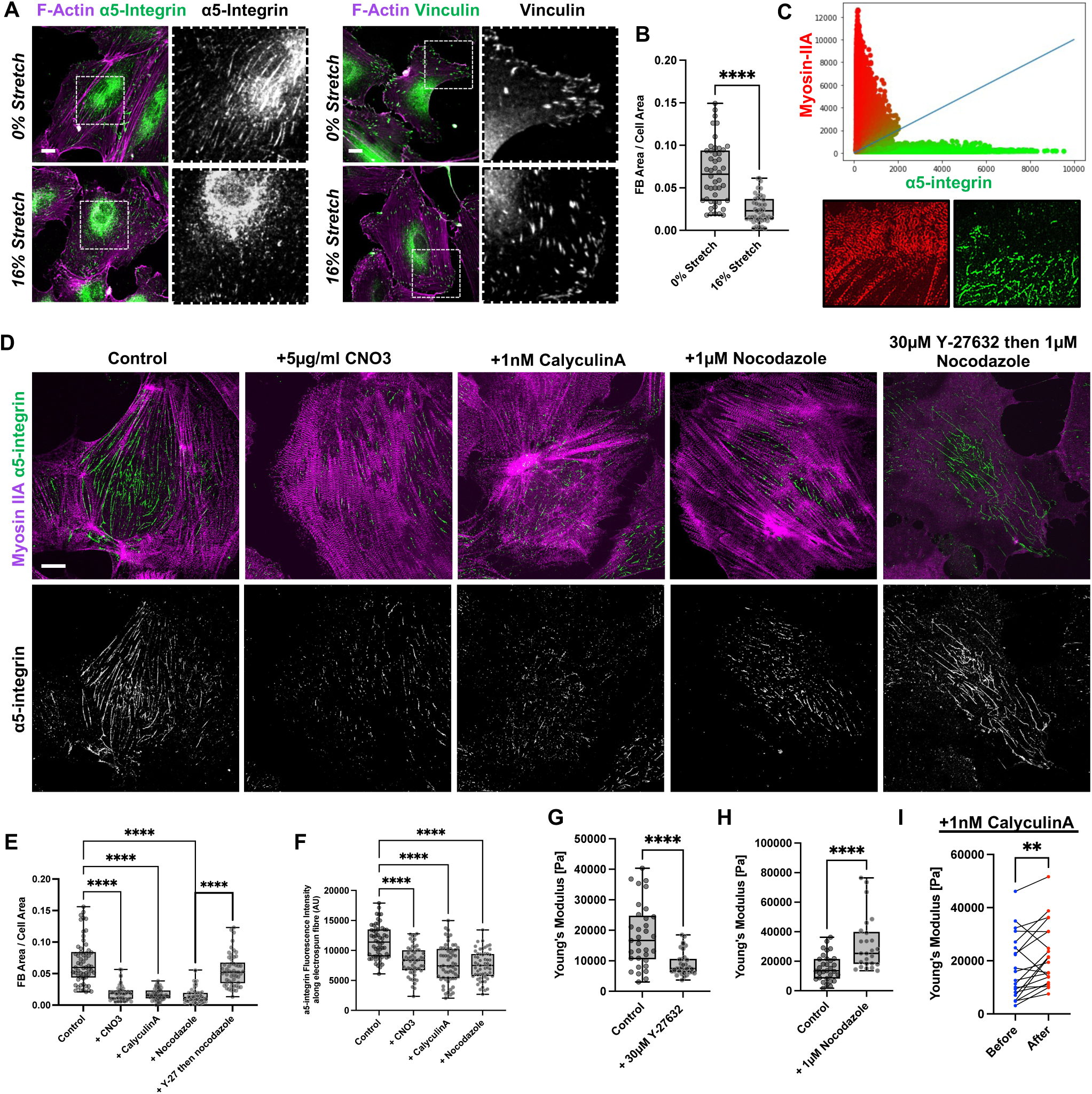
Fibrillar Adhesions are sensitive to myosin-II driven cortical stiffening. (A) HUVECs grown on plasma fibronectin-coated flexible PDMS in a stretching device, subjected to 0% stretch or a rapid 16% stretch and further incubation for 10 min. Cells visualized for F-actin (magenta) and α5-integrin (fibrillar adhesions, green, left) or vinculin (focal adhesions, green, right). Boxed areas shown with high magnification in a greyscale. Fibrillar adhesions decrease after substrate stretching while focal adhesions remain intact. Scale bar, 10µm. (B) Quantification of ratio of fibrillar adhesion area to total cell area (FB Area/Cell Area) for HUVEC under 0% stretch (*n*=46) or 16% stretch (*n*=47), *N*=3, box-and-whiskers plot (min, median, max) with all data points shown, *P*-values calculated using unpaired t-test, **** *P* < 0.0001. (C) Scatterplot showing pixel intensity correlations between myosin-IIA heavy chain (red) and α5-integrin (green), showing a strong relative mutual exclusivity (*n*=10 regions of interest from 10 cells). Representative images of myosin-IIA (red) and α5-integrin (green) in a region of interest of a HUVEC cell are shown. (D) HUVECs plated overnight on planar glass coverslips and stained for myosin-IIA heavy chain (magenta) and α5-integrin (green) under control conditions or after treatment with 5µg/ml Rho activator CNO3 for 3 hours, 1nM of MLCP inhibitor calyculin A (15 min) or 1 µM nocodazole (which increases Rho activity due to release of GEF-H1 upon microtubule disruption) for 1 hour. Note that activation of myosin filament formation results in disruption of fibrillar adhesions. Pre-treatment of cells with 30 µM Y-27632 for 15 min before co-addition of 1 µM nocodazole mostly preserves fibrillar adhesion formation. Scale bar, 10 µm. (E) Quantification of ratio of fibrillar adhesion area to total cell area (FB Area/Cell Area) for control HUVEC (*n*=58), cells treated with 5µg/ml CNO3 for 3 hours (*n*=51), 1nM calyculin A for 15 min (*n*=42), 10 µM nocodazole for 1 hour (*n*=59), or 30 µM Y-27632 added for 15 min before co-addition of 1 µM nocodazole for an additional hour (nocodazole and Y-37632) (*n*=54), *N*=3, box-and-whiskers plot (min, median, max) with all data points shown, one way ANOVA comparing means of + CNO3, + Calyculin A and + Nocodazole to control, as well as + Y-27 then Nocodazole to Nocodazole, **** *P* < 0.0001). (F) Mean intensity of α5-integrin clustering along electrospun nanofibers under control conditions (*n*=57) or after incubation with 5µg/ml CNO3 for 3 hours (*n*=56) or 1nM calyculin A for 15 min (*n*=60), or 10 µM nocodazole for 1 hour (*n*=57). (*N*=3, Box and whiskers (min, median, max) with all data points shown, *P*-values calculated using one way ANOVA, **** *P* < 0.0001. See supplementary Figure S5B for representative images. (G) Graph showing measurements of cortical stiffness (Pa) in HUVEC by atomic force microscopy (AFM) in control (*n*=33 cells) and cells pre-treated with 30µM Y-27632 for 1 hour (*n*=34 cells), *N*=3 independent experiments, box-and-whiskers plot (min, median, max) with all data points shown, two-tailed unpaired t-test, **** *P* < 0.0001. (H) Graph showing measurements of cortical stiffness (Pa) of HUVEC by AFM in control (*n*=31 cells) and cells pre-treated with 1µM nocodazole for 1 hour (*n*=25 cells) *N*=3 independent experiments, box-and-whiskers plot (min, median, max) with all data points shown, two-tailed unpaired t-test, **** *P* < 0.0001. (I) Matched cortical stiffness (Pa) AFM measurements in HUVEC before and after treatment with 1 nM calyculin A for 10–20 min (pooled data on *n*=20 cells, in *N*=2 independent experiments, paired t-test, ** *P* < 0.01).

Since cortical stiffness and tension are largely governed by actomyosin contractility [40, 41], we next tested whether increasing myosin-II activity would mimic these effects. We observed a striking mutual exclusivity between fibrillar adhesions and myosin-IIA filaments on planar substrates **(Fig. 5C)**. To directly assess the impact of myosin activation, we stimulated cells with the Rho activator CNO3, the myosin light chain phosphatase (MLCP) inhibitor calyculin A, and the microtubule disrupting drug nocodazole, which indirectly activates RhoA via GEF-H1 release from microtubules [42] **(Fig. 5D)**. All treatments led to disassembly of fibrillar adhesions and fibronectin fibrils **(Fig. 5D, E, S5A)** on planar glass. We confirmed that nocodazole inhibits fibrillar adhesion due to Rho activation and not loss of microtubules, as addition of ROCK inhibitor just before treatment preserved fibrillar adhesions **(Fig.5D, E).** Similarly, all three myosin-activating treatments (nocodazole, CNO3, and calyculin A) also reduced the formation of fibrillar adhesions along nanofibers **(Fig.5F, S5B)**. We also observed this by live-cell imaging of HUVECs co-expressing GFP-tensin3 and mCherry–myosin-IIA. Following nocodazole treatment, we observed rapid disassembly of fibrillar adhesions and increased focal adhesion formation, coinciding with the accumulation of adjacent myosin-IIA filaments **(Supplemental Movie3, Fig.S5C)**. To quantify the impact of these treatments on the cell cortex, we performed atomic force microscopy (AFM) using nano-indentation to probe the cortical regions (excluding the nucleus). Inhibition of myosin-II with Y-27632 led to a ∼50% decrease in cortical stiffness **(Fig.5G)**. Meanwhile nocodazole treatment doubled the cortical stiffness **(Fig.5H)**, while calyculin A increased cortical stiffness by ∼20% within 10-15 minutes in the same cell after treatment **(Fig.5I)**.

To augment membrane tension, we employed two established approaches: treatment with methyl-β-cyclodextrin (MβCD), which depletes membrane cholesterol [43, 44] and hypo-osmotic shock (1:3 medium dilution with distilled water) [39, 45]. Both treatments led to a marked reduction in fibrillar adhesions in HUVECs cultured on planar glass and nanofibers, while focal adhesions remained largely unaffected **(Fig.6A-D)**. To visualize this in real time, we transfected HUVEC with GFP-tensin3 on planar glass and subjected them to hypotonic conditions. We observed a rapid loss of GFP-tensin3 positive fibrillar adhesions from the central cell regions, whereas peripheral focal adhesions remained stable **(Supplemental Movie 4, Fig. 6E)**. We confirmed that membrane tension induced fibrillar adhesion disruption was not a result of Rho activation as addition of ROCK inhibitor before treatment with MβCD was unable to inhibit fibrillar adhesion disassembly **(Fig. S5C, D)**. These findings indicate that, in sharp contrast to focal adhesions which mature with increased actomyosin contractility, fibrillar adhesions are negatively regulated by excessive myosin-IIA filaments, cortical stiffening and elevated membrane tension.

**Figure 6.**
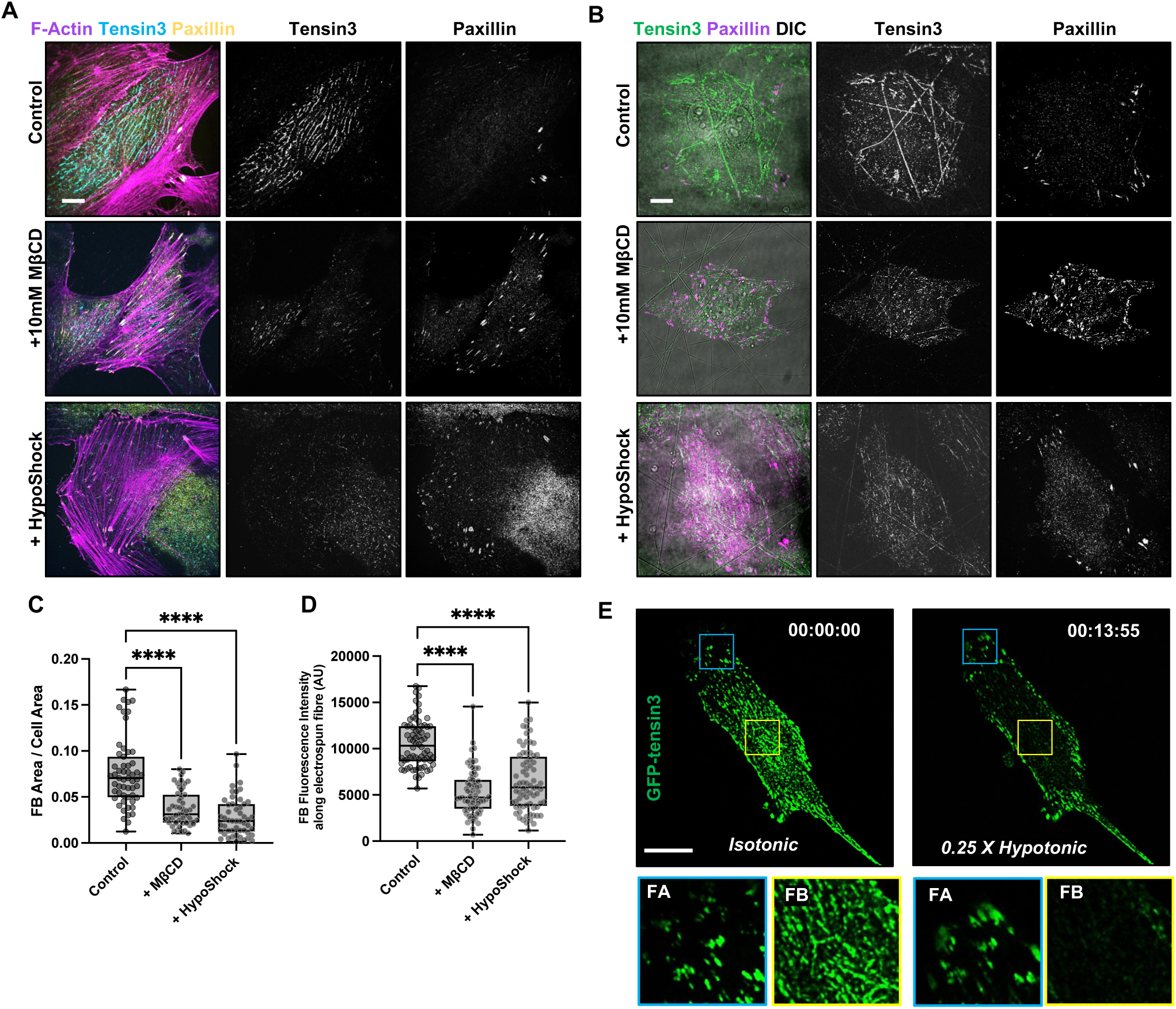
Fibrillar adhesions are sensitive to elevated membrane tension. (A) Cells spread on planar glass substrates under control conditions or in the presence 10 mM methyl-β-cyclodextrin (MβCD) for 90 minutes or hypo-osmotic medium (HypoShock – 0.25X hypotonic) for 20 minutes to increase membrane tension. Images show F-actin (magenta), fibrillar adhesions visualized by tensin3 (cyan) which disassemble, and focal adhesions visualized by paxillin (yellow) which are not affected. Scale bar, 10 µm. (B) Cells plated overnight on plasma fibronectin coated electrospun nanofibers under control conditions or treated with 10 mM MβCD for 90 min as well as hypo-osmotic medium (0.25X hypotonic) for 20 min, showing tensin3 (green) containing fibrillar adhesions assembled along nanofibers and paxillin (magenta) containing focal adhesions. Electrospun nanofibers shown in DIC (greyscale) in composite. Fibrillar adhesions are disassembled but focal adhesions are not affected upon MβCD and hypotonic medium treatment. Scale bar, 10 µm. (C) Quantification of ratio of fibrillar adhesion area to total cell area (FB Area/Cell Area) for control HUVEC (*n*=55), cells treated with 10 mM MβCD (*n*=51), and hypo-osmotic shock (0.25X hypotonic) (*n*=50), *N*=3, box-and-whiskers plot (min, median, max) with all data points shown, *P*-values calculated using one way ANOVA, **** *P* < 0.0001. (D) Mean intensity of fibrillar adhesions along electrospun nanofibers under control conditions (*n*=70), with the addition of 10 mM MβCD (*n*=71), or after treatment 0.25X hypotonic medium (*n*=72) (*N*=3, box-and-whiskers plot (min, median, max) with all data points shown, *P*-values calculated using one way ANOVA, **** *P* < 0.0001). (E) Still images from time-lapse imaging of HUVEC on planar glass transfected with GFP-tensin3 (green) under isotonic conditions and indicatable for 13:55 mins in 0.25x hypotonic medium. Boxed areas with peripheral focal adhesions (FAs, blue box) and central fibrillar adhesions (FB, yellow box) are shown at higher magnification in figures below. See Supplemental Movie 4. Fibrillar adhesions are disappearing while focal adhesions are not affected by hypo-osmotic shock. Scale bar, 10 µm.

### The clustering patterns of adhesion receptors across various topographical features can be explained by a unifying theoretical model

Our observations indicate that, in the presence of a topographical cue, the major determining factors for fibrillar adhesion formation are the mechanical properties of the cortex/membrane and the presence of α5β1-integrins. We developed a theoretical model based on our previous work [46] which accounts for planar substrates, and predicts clusters – organised as dots or lines patterns – that disappear with increased membrane tension. We now extend this model to curved substrates. We introduce the idealized “adhesion receptors” and assume that the free energy of these receptors is minimal when there is a tilt between the membrane and ligand-coated substrate planes (**Fig 7A)**. This disfavours the configuration of parallel membrane and substrates, a generic situation since the real adhesion receptors are hardly straight. The relevance of the corresponding free-energy terms therefore depends on the actual shape or geometry. The two-spring representation in Figure 7A provides a particularly relevant physical picture for α5β1 integrin, as explained further in the Discussion.

**Figure 7.**
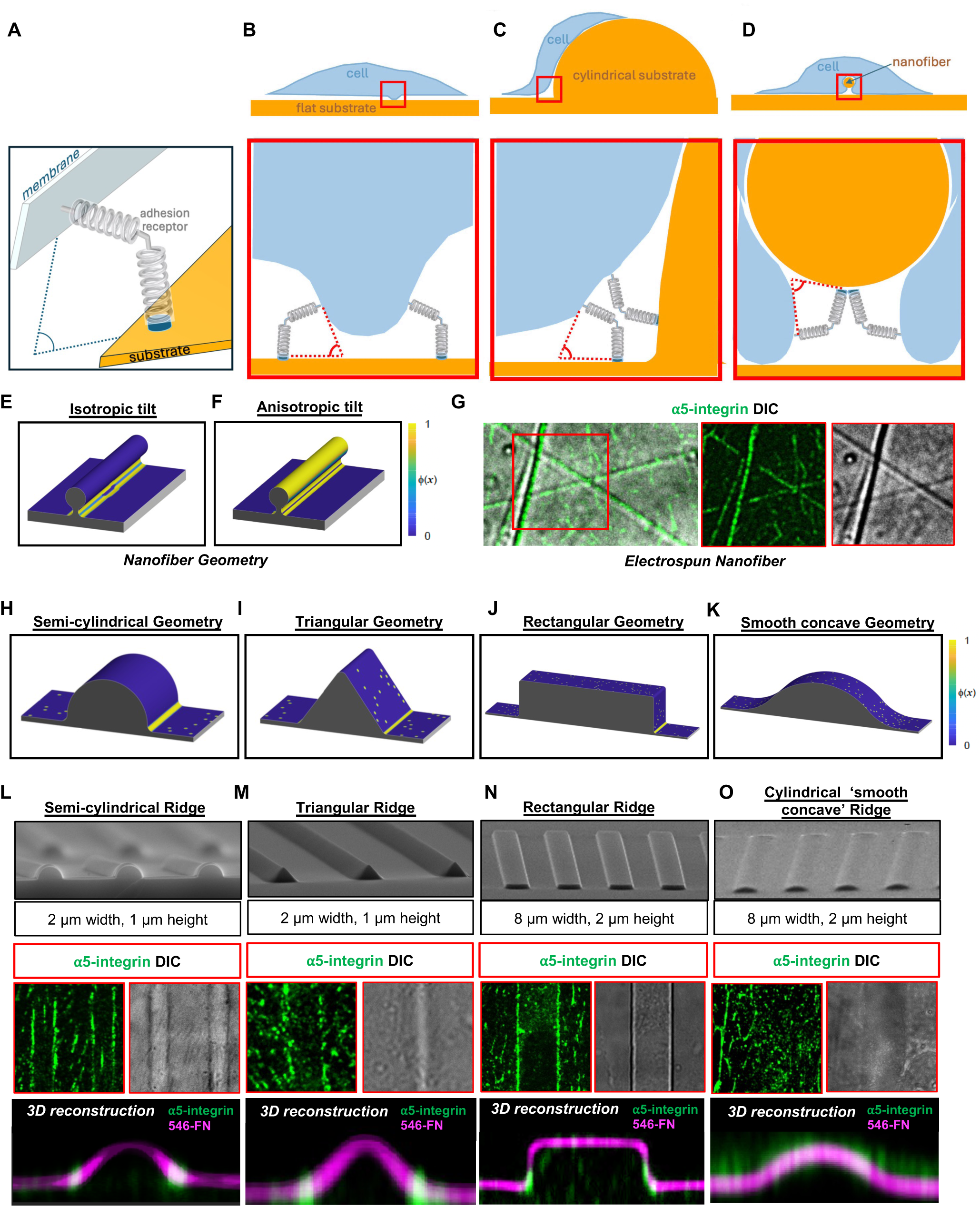
The clustering patterns of adhesion receptors across various topographical features can be explained by a unifying theoretical model. (A) The adhesion receptor is depicted as two springs connected at an angle to each other. The main assumption of the theory is that free energy approaching minimum when the plane of the cell membrane from which the receptor is protruding (grey) and plane of the substrate covered with the receptor’s ligand (orange) are tilted relatively to each other. (B, C, D) Situations when the tilted configuration can be approached. Upper row: entire cell (blue) interacting with the substrate (orange); the areas where adhesions are preferentially formed are boxed (red). Lower row: boxed areas at high magnification, showing the positions of adhesion receptors (not in scale) and the angles between the local tangents to cell membrane and the substrate (red dotted lines). (B) Cell on the flat substrate forming a small protrusion/fold (this situation was analyzed in detail in [46]). (C) Cell spreading over the semi-cylindrical pattern (cross-section). The adhesion receptors are concentrated along the concave edges of the semi-cylinders forming grooves with high negative curvature. (D) Cell engulfing a cylindrical nanofiber with a small diameter and high positive curvature. The adhesion receptors are concentrated along the nanofiber. (E, F) Numerical calculation of the clustering pattern of adhesion receptors along the nanofiber geometry calculated based on the assumption of (E) an isotropic receptor tilt and (F) an anisotropic receptor tilt. Fraction of adherent receptors Φ(*x*) represented by spectrum scale on the right. (G) (left) SIM merged image of α5-integrin (green) clustering along electrospun nanofibers (DIC, grey) in HUVEC. The boxed area is shown in individual channels in images on the right. (H-K) Numerical calculation of clustering patterns of adhesion receptors (based on the assumption of isotropic receptor tilt) on various curved geometries. (H) a semi-cylinder geometry with concave edges of high negative curvature (I) a triangle geometry with concave and convex edges, (J) a rectangle geometry with concave edges, and (K) a geometry with smooth concave edges. Calculated fraction of adherent receptors Φ(*x*) is represented by spectrum scale on the right. Note that linear receptor clustering occurs preferentially along concave edges with high negative curvature. (L-O) Experimental validation of numerical calculations in (H-K). Top panels show SEM images of PDMS microfabricated substrates of (L) semi-cylindrical substrates with concave edges, (M) triangular substrates with concave and convex edges, (N) rectangular substrates with concave edges, and (O) cylindrical segments with smooth concave edges. Middle panels are 2D confocal images of HUVEC plated on corresponding plasma fibronectin-coated PDMS patterns showing α5-integrin (green) localization along the micro-ridge edges (DIC, grey). The bottom panels show confocal 3D reconstructions of α5-integrin (green) clustering along the respective micro-ridge geometries with plasma fibronectin coating labelled using fluorescent fibrinogen (magenta) to visualize the ridge surface.

Intuitively, the assumption about the preferential tilt explains that either membrane protrusion **(Fig 7B)** or substrate topography containing ridges with concave edges of large negative curvature will promote the receptor clustering **(Fig 7C).** Moreover partially engulfed nanofibers with high positive curvature contain regions where tilt-induced clustering of receptors can occur **(Fig 7D)**. Compared with flat substrates, concave geometries or the nanofibers reduce the mechanical deformation energy of the membrane, thereby facilitating formation or receptor clusters.

In more detail, we described a membrane-protein-substrate system by two fields: (1) the distance between the membrane and the substrate, denoted *e*(***x***) with ***x*** being the spatial coordinate of the curved substrate; (2) the fraction of bound cell adhesion receptors, denoted *ϕ*(***x***) ∈ (0, 1). The stable clustering pattern of cell adhesion receptors is dictated by minimizing the free energy of the system, denoted *F*. Following our previous work [46], we considered four contributions to the free energy of the system,

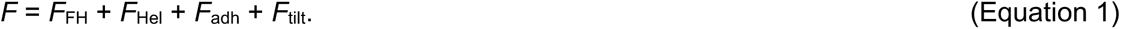

Here, the first term *F*_FH_ describes the mixing of cell adhesion receptors with ligands (binders). The second term *F*_Hel_ represents the mechanical deformation of the membrane (including two parts, i.e., membrane tension and bending). The third term *F*_adh_ corresponds to the adhesion interactions between the membrane and the substrate. The last term *F*_tilt_ quantifies the tilt effect of cell adhesion receptors **(Fig. 7A)**. The equations corresponding to each 4 terms are shown in **Box_1** and explained in detail in Supplemental Theory material. Note that *F*_tilt_ has an anisotropic curvature dependent term. Indeed, if like in Figure 7A the substrate shape is invariant in a given direction (orthogonal to the figure plane) in this direction the gain in energy due to intrinsic receptor tilt does not take place. The curvature-dependent term accounts for the effect of 90° rotation of the adhesion receptor around its axis (see the equation for *F*_tilt_ in **Box_1** and Supplementary Theory material).

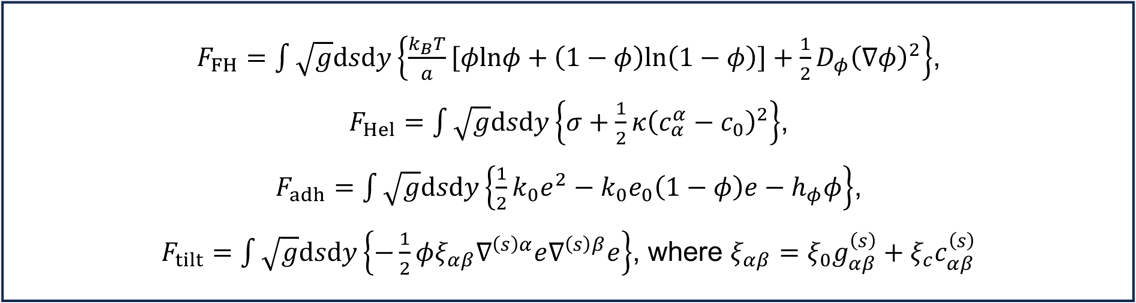

In this box, we give the detailed expressions of *F*_FH_, *F*_Hel_, *F*_adh_, and *F*_tilt_, where *F*_FH_ is expressed as in the Flory–Huggins theory; *F*_Hel_ is expressed in the classical Helfrich formalism; *F*_adh_ includes a spring interaction part and chemical potential part. *F*_tilt_ is expressed in a generic case where the protein tilt effect can be anisotropic and depends on the substrate curvature. See the Supplementary Theory material for detailed explanation for each term.

We performed numerical calculations with substrate geometries exhibiting modulated curvature along only one direction, namely along the *x*-axis, and invariant along the *y*-axis **(Fig S6A-F)**. We first applied our model to the electrospun nanofiber geometry and found it correctly recapitulated the α5β1-integrin adhesion – continuous clustering along the nanofiber observed experimentally (**Fig. 7G**), in sharp contrasts to flat substrates, where no such continuous clustering occurs (**Fig. 1A**, and [46]). This clustering was predominantly at the fiber base, indicating to partial engulfment of the nanofibers **(Fig. 7E)** as depicted in **Fig. 7D**. Limitations in microscopic resolution prevented precise determination of whether α5β1-clusters localized strictly beneath, partially wrapped, or fully encircled the fiber. However, the assumption that tilt between the cell membrane and substrate surface should be anisotropic and depend on surface curvature leads to prediction that adhesion receptor clustering occurs along the entire surface of the fiber (**Fig. 7F**, see Discussion and Supplementary Theory material for more details).

We next examined how the adhesion receptor with intrinsic tilt would respond to substrates with ridges of different geometries. Using our numerical calculations (see Supplementary Theory material for more details), we find that what is critical is not the overall shape of the ridges (e.g., semi-cylindrical, triangular, rectangular shapes), but the existence of concave edges with sufficiently high negative curvature to ensure the formation of a large-scale, continuous, receptor clusters **(Fig. 7H-J)**; so that no large-scale linear clustering would occur along smooth concave edges **(Fig. 7K)**.

To validate these predictions experimentally, we fabricated polydimethylsiloxane (PDMS) substrates containing microarrays of parallel ridges with the same geometries used in our numerical calculations **(Fig. 7L-O, S7A)**. When HUVECs were cultured on ridges with semi-circular, triangular, and rectangular profiles with the edges with high negative curvature, α5-integrin consistently formed two parallel lines flanking either side of each ridge. Importantly, 3D reconstruction of our confocal images revealed that α5-integrin clusters specifically localized at the concave but not convex edges of the features **(Fig.7L-N)**, precisely as predicted by the model **(Fig.7H-J)**. In contrast, cells plated on cylindrical ridges with smooth concave edges did not exhibit localized α5-integrin clustering **(Fig.7O)**, in further agreement with our computation **(Fig.7K)**. We confirmed that clustering behaviour was specific to fibrillar adhesions, as paxillin positive focal adhesions showed no such curvature-dependent localization **(Fig. S7B)**. Furthermore, fibrillar adhesion formation along concave regions was still observed when cells were pre-treated with ROCK inhibitor prior to plating **(Fig. S7C)**.

These results suggest that adhesion receptor clustering along specific topographical features can be explained by their tilt preferences. Evidence that α5β1-integrin behaves similarly to these theoretically defined adhesion receptors will be presented in the Discussion.

## Discussion

Our results suggest a new paradigm in the understanding of fibrillar adhesion formation. Contrary to the long-standing model that fibrillar adhesions arise from myosin-II–dependent segregation of mature focal adhesions [12], we demonstrate that fibrillar adhesions form independently, in direct response to local membrane curvature, guided by nanoscale substrate topography. This distinction became apparent when comparing planar substrates with those containing fibres or concave edges with high negative curvature. On planar substrates, where fibrillar adhesions were originally described, their formation requires fibronectin secretion, tensins, and myosin-IIA activity. However, when cells are cultured on decellularized ECM or electrospun nanofibers, these requirements become dispensable. Notably, focal adhesions fail to align or respond to these topographical features, further supporting the idea that fibrillar adhesions represent a mechanistically distinct system, specialized for matrix topography recognition.

Dissection of the relationship between fibrillar adhesion and fibronectin fibrillogenesis has been difficult due to their interdependence on planar substrates. Introducing substrates with nanotopographical features overcomes this limitation. While the formation of fibrillar adhesions along nanofibers was independent of myosin activity, fibronectin fibrillogenesis is strictly myosin-dependent. Of note, upon ROCK inhibitor washout, fibronectin fibrils rapidly reassemble in close proximity to pre-existing fibrillar adhesions, suggesting a positive feedback mechanism in which fibrillar adhesions induce fibronectin fibrillogenesis, promoting formation of cell-derived nanofibers, which in turn reinforces fibrillar adhesion formation via curvature-mediated assembly. This feedback could be especially important for formation of fibrillar adhesions on the flat substrate. However, it remains unclear whether myosin’s role in fibronectin fibrillogenesis primarily involves the regulation of fibronectin secretion [18], the mechanical unfolding of cryptic fibronectin-binding sites [19–22], or local modulation of cell cortex [41]. It is likely that all these factors are working in concert.

Response of cells to local changes of membrane curvature induced by external topographical features in some cases is mediated by accumulation of BAR domain proteins in the regions of higher curvature. In particular, in recently discovered “curved adhesions” a direct interaction between β5-integrin and the F-BAR domain protein FCHo2 was detected [10]. Unlike the curved adhesions, we found that fibrillar adhesions contain neither β5-integrin, nor FCHo2, nor several other BAR domain proteins we examined.

Rapidly accumulating data shows that processes of liquid-liquid phase separation (LLPS) could be involved in self-organization of the membrane associated proteins. In particular several proteins associated with integrin adhesions were shown to undergo LLPS including talin [47], p130CAS [48], KANK1 [32], tensin3 [35] and tensin1 [33, 34]. The possible role of tensin LLPS in the assembly of fibrillar adhesions is especially interesting since these proteins are major components of these structures. However, we have shown that formation of α5β1-integrin clusters along nanofibers occurs even in tensin1&3 knockdown cells. Moreover tensin can be extracted from the fibrillar adhesions by the solvent 1,6-Hexanediol, leaving the α5-integrin clusters apparently intact. Thus, formation of fibrillar adhesions should be explained based on specific features of α5β1-integrin, the primary integrin found in these adhesions, and its relationship with the membrane.

In stark contrast to focal adhesions that grow in response to myosin-II driven force [49, 50], here we show that fibrillar adhesion disassemble upon myosin-II activation by elevation of RhoA (CNO3, nocodazole) or inhibition of MLCP (Calyculin A). Prior studies have also demonstrated that nocodazole treatment disrupts fibrillar adhesions [51] and calyculin A blocks fibronectin fibrillogenesis [20]. Our AFM data shows that both nocodazole and calyculin A treatment induce cortical stiffening, whereas ROCK inhibition by Y-27632 reduces it. Consistently, our data suggests that fibrillar adhesions in cells treated with ROCK inhibitor exhibit a somewhat enhanced association to pre-existing CDM and electrospun nanofibers compared to untreated controls. This indicates that a softer cell cortex facilitates the formation of fibrillar adhesions along topographical features. Accumulating data highlights the tight relationship between myosin-II dependent cortical tension and plasma membrane tension [40, 41, 52, 53]. Consistently our data demonstrates that fibrillar adhesion disassembly is also triggered by treatments that increase membrane tension. Of note, this regulatory profile parallels that of podosomes, which disassemble upon elevated membrane tension and myosin overactivation, and also preferentially assemble along micro-topographical features such as triangular micro-ridges [39].

Altogether our data suggests that external topographical features inducing high curvature of the cell membrane facilitate formation of α5β1-integrin clusters while increase of cortical/membrane tension disrupts receptor clustering. We proposed a theory, where adhesion receptor clustering is seen as a sort of wetting-dewetting transition [54, 55] including the assumption that the free energy of the adhesion receptor is minimized when the cell membrane plane and ligand-coated substrate plane are tilted relative to each other **(Fig 7A)** [46]. This theory predicted the pattern of adhesion receptor clusters on the planar substrate, as well as the negative effect of membrane tension on receptor cluster formation [46]. Both these predictions were confirmed in our experiments. Further, we expanded this theory to the situation when the substrate is non-planar but has specific curved topographical features **(see Fig 7 and Supplemental Theory material)**. The theory predicts and experiments confirm that concave edges of the micropatterned ridges with high negative curvature are most preferable sites for the clustering of adhesion receptors. The theory also accounts for the observed preferential clustering of the receptors in the concave cleft between the nanofiber and the substrate. The additional assumption of anisotropy of adhesion receptor tilt (equation for *F*_tilt_ term) makes it possible that not only this concave region but the ‘top’ convex strip on the nanofiber surface could be the region of preferential clustering of adhesion receptors. This can be inferred from the assumption that the substrate curvature can play a role in the tilt orientation. Our numerical calculations showed that the entire surface of the fiber will be coated with the adhesion receptors if the tilt effect is enhanced at concave regions while suppressed at convex regions (See further details in Supplementary Theory material). Such an anisotropic tilt assumption does not change the experimentally verified theoretical predictions of receptor clustering at concave edges of micropatterns **(Fig. S8)**. Localization of integrin clustering on the top or/and beneath the nanofibers requires additional elucidation. Importantly, both versions of the theory account for experimentally observed preferential clustering of adhesion receptors along the topographical features as compared to planar surfaces.

What are the relationships between the theoretically defined adhesion receptor we introduce in this model and real α5β1-integrin which is the only integrin involved in the formation of fibrillar adhesions? First, the conformation of bent (“closed” or “resting”) α5β1 resembles the configuration of adhesion receptor in our model, namely, in principle, bent α5β1 fits the position when membrane and substrate planes are tilted relatively to each other **(Fig 7A,B)**. Indeed, fibrillar adhesions are enriched in β1-integrin in its bent conformation recognized by mAb13 antibody. Moreover, only α5β1 integrin, but not other integrin types that can bind RGD, has the appropriate shape in bent conformation suitable to connect the membrane and ligand coated substrate tilted relatively to each other: extracellular portion of α5β1 bent approximately 90 degrees, while other types of RDG binding integrins are bent 180 degrees, see Figure S9A [56, 57]. This may explain the exceptional role of α5β1 in formation of fibrillar adhesions. Even though bent conformation of α5β1 has lower affinity to fibronectin than extended conformation, it still can bind fibronectin, even faster than the extended conformation and is more abundant at the cell surface [58]. Moreover, upon binding the ligand, this integrin undergoes very rapid transition to extended open conformation with high affinity, bypassing the “extended-closed” state [59].

Thus, we hypothesize that α5β1-integrin forms clusters along topographical features or membrane folds (as predicted by our theory) still in its bent conformation **(Fig. S9B**). Relatively low affinity of the receptors at this conformation could facilitate the selective accumulation of these receptors in the regions of spontaneous or substrate-induced membrane bending. Further stages in assembly of fibrillar adhesions including integrin extension and activation (**Fig. S9C)** recruitment of tensin and its possible phase separation [34, 35], and formation of links with sub-membranous actin networks are not considered by this theory and full understanding of this process requires further studies. It is also worth noting that some integrins in extended conformation (such as αvβ3) could be also tilted relative to the cell membrane [5, 60], however this observation is hardly applicable to the phenomenon of formation of fibrillar adhesions since it does not explain the specific role of α5β1-integrin.

In conclusion, our study shows that fibrillar adhesions provide a mechanism for recognition of special topographical features by cells, specifically fibers, grooves or ridges with high cross-section curvature. This mechanism is based on tilt-induced and curvature-dependant phase separation of adhesion receptors and does not require BAR domain proteins. Secretion of cellular fibronectin, tensins and myosin-II may promote fibrillar adhesion formation on planar substrates, while increased membrane/cortical tension leads to disassembly of these structures. A physical model based on the assumption that adhesion receptors prefer to bind the ligand-coated surface where this surface is tilted relatively to the membrane plane comprises the first principle-based explanation of the receptor phase separation along topographical features and consistent with the specific role of α5β1, but not other integrins, in this process. Fibrillar adhesions may represent a rapid, topography driven form of cell adhesion, particularly relevant in 3D environments.

## Supporting information

Supplemental Theory

## Supplemental Figures

**Figure S1.**
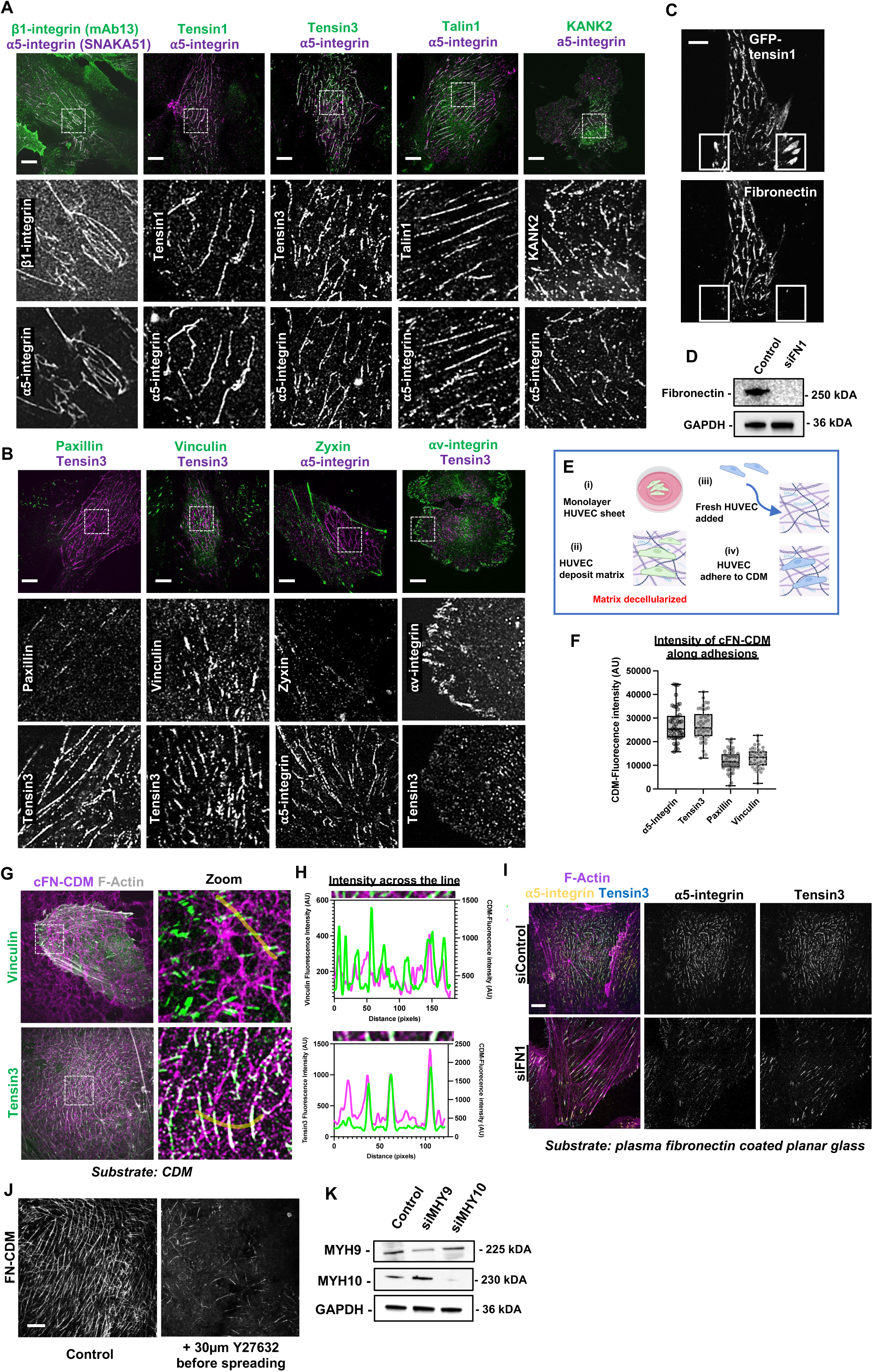
Fibronectin secretion and actomyosin contractility are required for fibrillar adhesion formation on planar substrates but dispensable on preformed cell-derived matrix. (A) Images of cells plated on planar glass stained for α5-integrin in its open conformation (SNAKA51, magenta) together with the bent conformation of β1-integrin (MAB13, green) on the left column. Other columns show co-localization of α5-integrin with other reported fibrillar adhesion components (green: tensin1, tensin3, talin1, KANK2). Greyscale images in middle and bottom rows show individual channel images of boxed regions. Scale bar, 10µm. (B) Images comparing localization of fibrillar adhesion components (tensin3 and α5-integrin/SNAKA51) along with focal adhesion components on planar glass. Columns from left to right show paxillin with tensin3, vinculin with tensin3, zyxin with α5-integrin, and αv-integrin with tensin3. Greyscale images in middle and bottom rows show individual channel images of boxed regions. Scale bar, 10µm. (C) HUVEC transfected with GFP–tensin1 and plated on planar glass coverslips. GFP-tensin1 displays two distinct localizations: elongated central fibrillar adhesions and peripheral focal adhesion plaques. Immunostaining for fibronectin reveals a co-localization with fibrillar adhesions and a lack of co-localization along focal adhesion sites (boxed). Scale bar,10 µm (D) Western blot showing FN1 knockdown efficiency in FN1 RNAi depleted cells compared to RNAi control using an anti-fibronectin antibody and GAPDH as a loading control. Respective molecular weights indicated. (E) Schematic illustration of cell derived matrix (CDM) preparation using HUVEC. Experimental setup adapted from [30]. (F) Quantification of the mean fluorescence intensity (AU) of the underlying cellular fibronectin in the pre-existing CDMs (cFN-CDM) along specific adhesions (Fibrillar adhesions: α5-Integrin *n*=45, tensin3 *n*=42; Focal adhesions: paxillin *n*=44, vinculin *n*=41). Box-and-whiskers-plot (min, median, max) with all data points shown. (G) HUVEC plated on pre-existing CDM and stained for cellular fibronectin to visualize the CDM (cFN-CDM, magenta), F-actin (phalloidin, grey), and vinculin (green, upper row) or tensin3 (green, bottom row). Scale bar, 10 µm. Images in the right column show higher magnification of the boxed areas in the left column. The line scans along yellow lines are shown in H. (H) Line scan along the yellow lines indicated in G, plots the fluorescence intensities for vinculin and tensin3 (green), together with corresponding underlying cellular fibronectin (cFN marking CDM, in magenta). (I) Cells grown on human plasma fibronectin coated planar glass coverslips, showing F-actin (magenta), α5-integrin (SNAKA51, yellow), and tensin3 (cyan), in siControl or after FN1 knockdown (siFN1). Loss of fibronectin secretion results in re-localization of α5-integrin and tensin3 to focal adhesion sites at the termini of stress fibers. Scale bar, 10 µm. (J) Western blot showing knockdown efficiency of myosin-IIA (siMYH9) and myosin-IIB (siMYH10) in HUVEC, using antibodies against MYH9 and MHY10, with GAPDH as loading control. Respective molecular weights indicated. (K) Images of cellular fibronectin in CDMs (cFN-CDM) secreted after 24 hours in control HUVEC or under treatment with 30µM Y-27632. Images showing failure to assemble fibronectin fibrils under ROCK inhibition. Scale bar, 10 µm.

**Figure S2.**
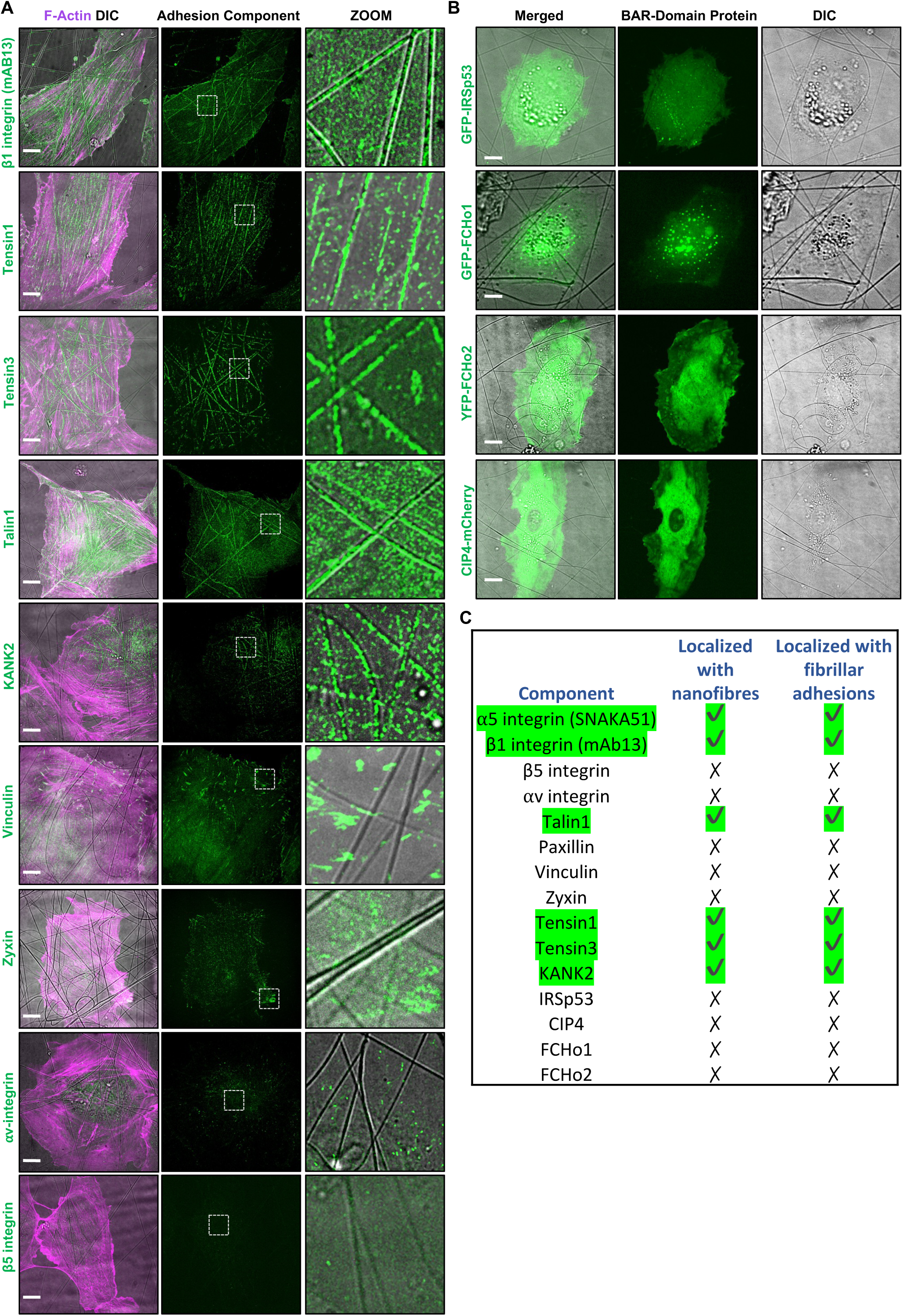
Fibrillar adhesion components associate with electrospun nanofibers. (A) SIM images of cells plated on plasma fibronectin–coated electrospun nanofibers (DIC, grey), with F-Actin (magenta) and multiple adhesion components shown in green. Images in the right column show zoomed boxed areas merged with DIC nanofiber images. Scale bar,10 µm. (B) Images of HUVEC transfected with fluorescently tagged BAR-domain proteins: I-BAR (GFP-IRSP53) and F-BAR (mCherry-CIP4, GFP-FCHo1, YFP-FCHo2) plated overnight on plasma fibronectin–coated electrospun nanofibers. No enrichment was observed along nanofibers (seen in DIC, grey). Scale bar, 10µm. (C) Table summarizing adhesion or BAR-domain protein localization along electrospun nanofibers and fibrillar adhesions on planar substrates.

**Figure S3.**
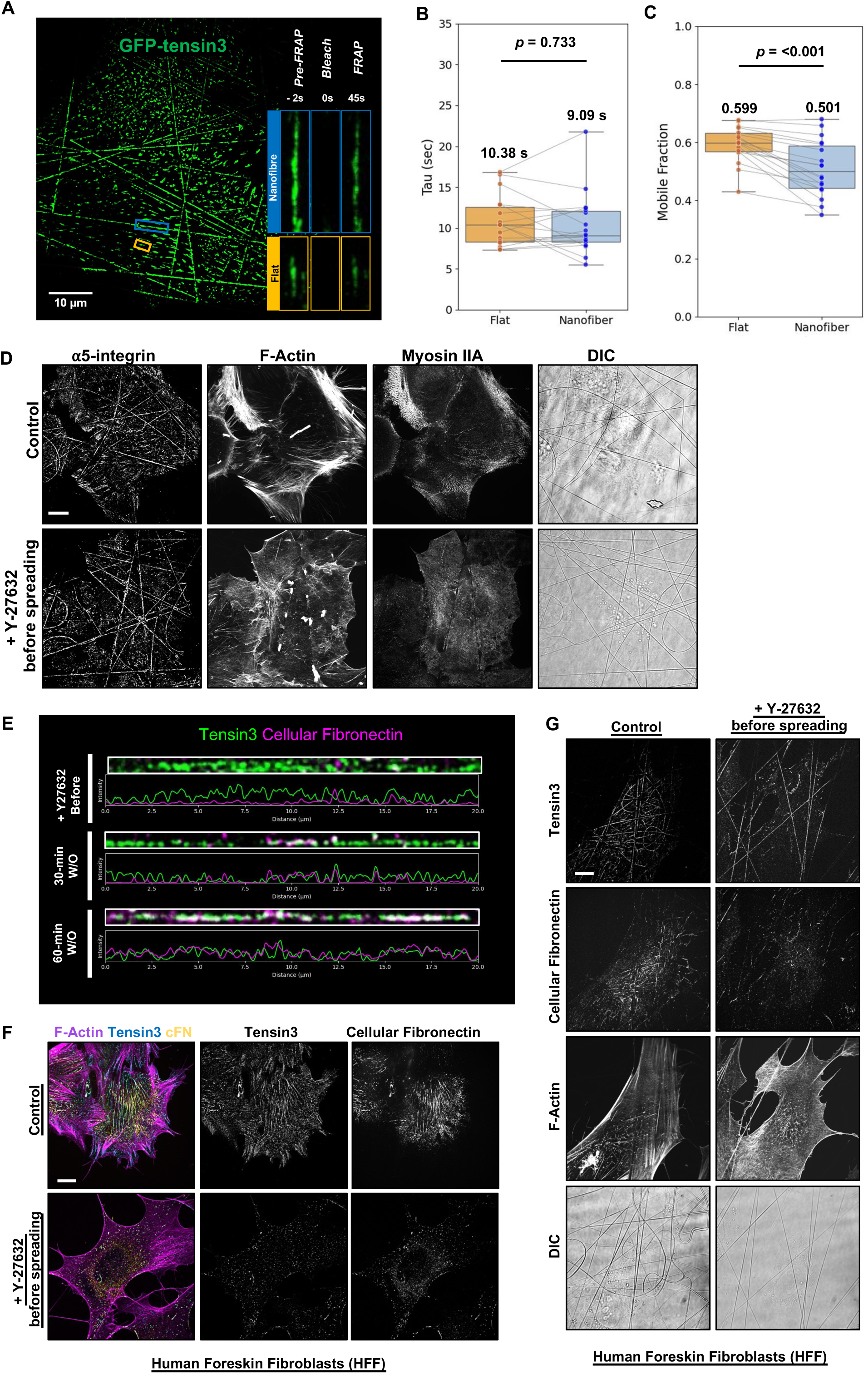
Formation of fibrillar adhesions along topographic cues is independent of actomyosin and fibronectin secretion. (A) Image of HUVEC transfected with GFP–tensin3 and plated on human plasma-fibronectin coated electrospun nanofibers used for evaluation of fibrillar adhesion dynamics on flat versus nanofiber regions by FRAP. Insets show adhesions formed in flat (orange box) or nanofiber (blue box) regions at the state of pre-bleaching (−2s), photobleaching (0), and recovery (45s). Scale bar, 10µm. See also Supplemental Movie 2. (B) Graph showing time recovery constants determined by single exponential fit (tau τ, seconds) of GFP-tensin3 fibrillar adhesions on flat versus nanofiber regions, with median values indicated on the graph (*n*=18, *N*=3, box-and-whiskers plot (min, median, max) with all data points shown, *P*-values calculated using Wilcoxon signed-rank test) (C) Graph showing mobile fractions of GFP-tensin3 fibrillar adhesions on flat versus nanofiber regions determined by single exponential fit, with median values indicated on the graph (*n*=18, *N*=3, box-and-whiskers plot (min, median, max) with all data points shown, *P*-values calculated using Wilcoxon signed-rank test) (D) HUVEC plated on plasma fibronectin coated electrospun nanofibers under control conditions or under treatment with 30µM ROCK inhibitor Y-27632 and stained (from left to right) for α5-integrin, F-actin and myosin-IIA heavy chain. Corresponding DIC images showing nanofibers are in the right column. Scale bar, 10µm. (E) Representative images of distribution of tensin3 (green) and cellular fibronectin (magenta) together with line scan intensity profiles along electrospun nanofibers in cells plated in the presence of 30 µM Y-27632 (upper panel), and after washing out the drug for 30 minutes (middle panel) or 60 minutes (lower panel). Accumulation of fibronectin after ROCK inhibitor washout correlates with the intensity of tensin3 staining. (F) Human foreskin fibroblasts (HFF) cultured overnight on planar glass in control conditions or with 30µM Y-27632 added before spreading. Left columns show merged images of F-actin (magenta), tensin3 (cyan), and cellular fibronectin (cFN; yellow). Individual grayscale panels for tensin3 and cellular fibronectin are shown in the middle and right columns. Scale bar, 10µm. (G) Images showing tensin3, cellular fibronectin and F-actin in HFFs plated on human plasma fibronectin coated electrospun nanofibers (DIC) under control conditions or in the presence of 30 µM Y-27632. Like HUVEC, fibrillar adhesions in HFF also form along nanofibers under ROCK inhibition, however formation of fibronectin fibrils is reduced. Scale bar, 10µm.

**Figure S4.**
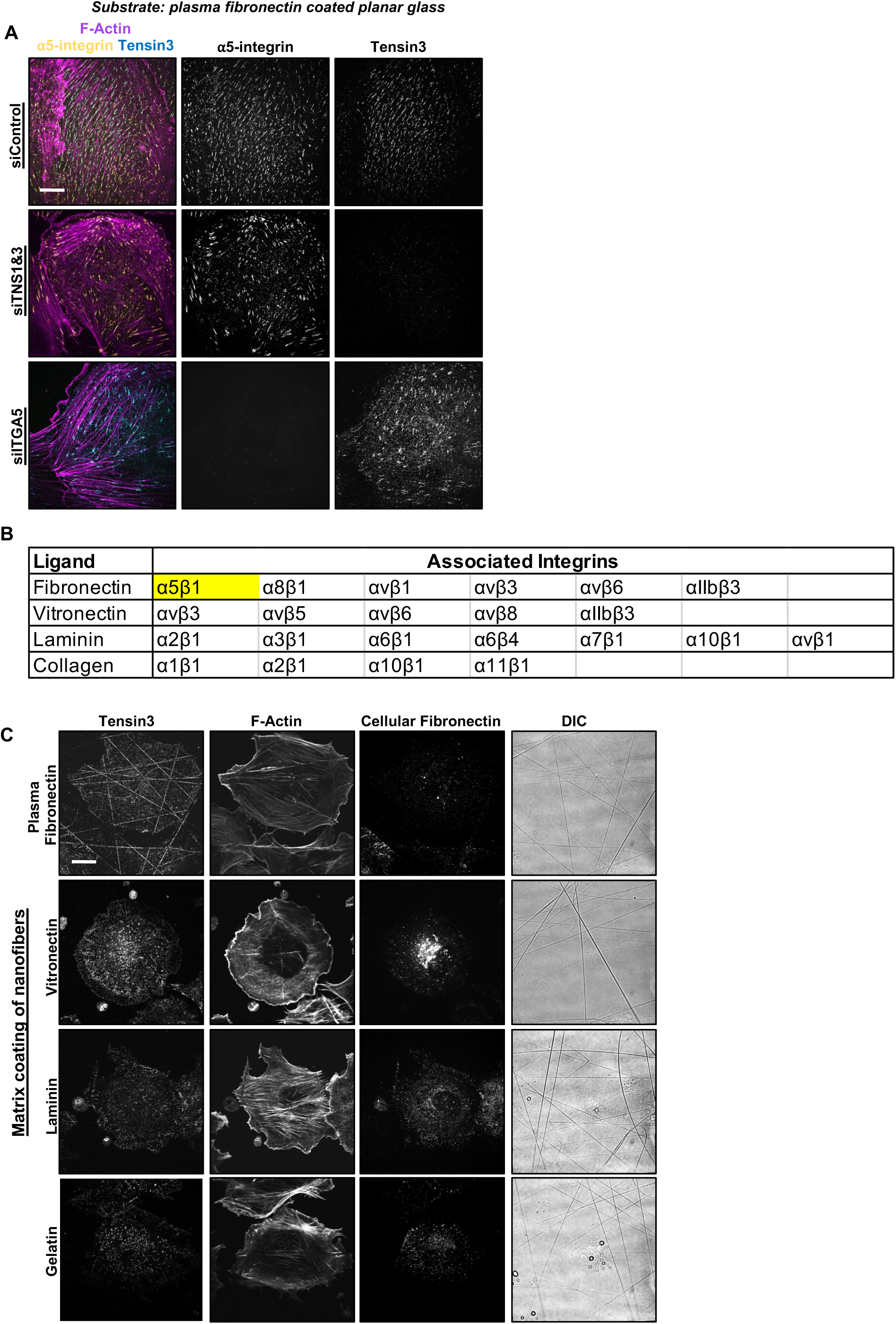
Formation of α5-integrin clusters along electrospun nanofibers is independent of tensins. (A) Images of HUVEC plated on human plasma fibronectin-coated planar glass coverslips, stained for F-actin (magenta), α5-integrin (yellow), and tensin3 (cyan). Left column shows merged images representing cells under control conditions (siControl), upon double knockdown of tensin1 and tensin3 (siTNS1&3), and knockdown of α5-integrin (siITGA5). Single channel greyscale images of the same cells are shown in in columns in the middle and right. Both knockdowns result in disappearance of fibrillar adhesions and re-localization of remaining adhesion components to focal adhesion sites at the termini of F-Actin stress fibers. (The adhesion of these cells to nanofibers is shown in Fig. 4D). Scale bar, 10 µm. (B) Table listing the integrin receptors associated with different ECM ligands: fibronectin, vitronectin, laminin and collagen (adapted from [61]). (C) Images of cells fixed 30 minutes after plating on electrospun nanofibers coated with either: 10 µg/mL human plasma fibronectin, 20 µg/mL vitronectin, 10 µg/mL laminin or 0.1% gelatin. Immunofluorescent staining for tensin3, cellular fibronectin and phalloidin staining of F-actin along with DIC images of electrospun nanofibers are shown. Only coating of the substrate with human plasma fibronectin enables the formation of tensin3 positive fibrillar adhesions along electrospun nanofibers. Scale bar, 10 µm.

**Figure S5.**
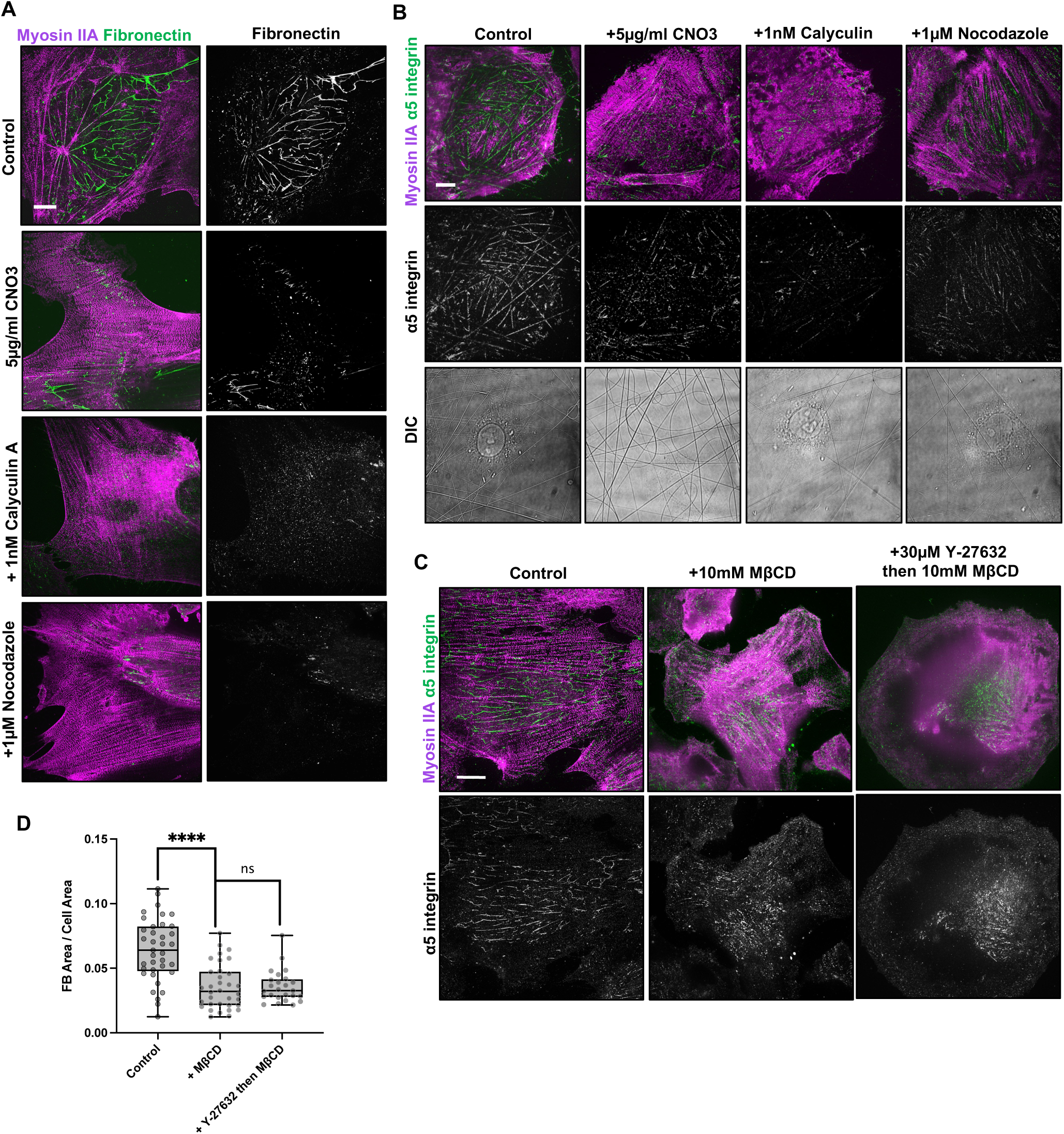
Fibrillar adhesions are sensitive to myosin-II overactivation and increased membrane tension. (A) HUVECs grown on planar glass coverslips. Left column shows cells stained for myosin-IIA heavy chain (magenta) and fibronectin (green) under control conditions and after treatment with 5µg/ml Rho activator CNO3 for 3 hours, 1nM of MLCP inhibitor calyculin A for 15 min and 1µM nocodazole (activating RhoA via GEF-H1 release following microtubule depolymerization) for 1 hour. The right column shows greyscale images of fibronectin fibers. Increase in myosin-IIA filaments results in disappearance of fibronectin fibrils. Scale bar, 10 µm. (B) Cells grown on electrospun nanofibers under control conditions and after treatment with 5µg/ml CNO3 for 3 hours, 1nM calyculin A for 15 min or 1µM nocodazole for 1 hour which all promote myosin filament formation. Top rows show merged images of myosin-IIA (magenta) and α5-integrin (green). Greyscale images show α5-integrin in corresponding cells (middle row) and nanofibers visualized by DIC (bottom row). Increase in myosin-IIA filaments disrupts clustering of α5-integrin along nanofibers. Scale bar, 10µm. (C) Cells plated overnight on planar glass, under control conditions (left column), treatment with 10 mM methyl-β-cyclodextrin (MβCD) for 90 mins (central column), or pre-treatment with 30 µM Y-27632 for 15 min and co-addition of 10 mM MβCD for 90 mins. Top row shows merged images of myosin-IIA (magenta) and α5-integrin (green), bottom row shows greyscale images of α5-integrin in corresponding cells. Pre-treatment of cells with ROCK inhibitor before co-addition of MβCD does not preserve fibrillar adhesions. Scale bar, 10 µm. (D) Quantification of ratio of fibrillar adhesion area to total cell area (FB Area/Cell Area) for cells under control conditions (*n*=37), treated with 10 mM MβCD (*n*=34), or pre-treated with 30 µM Y-27632 followed by co-addition of 10 mM MβCD (*n*=25). *P*-values calculated using one way ANOVA, ns, not significant, **** *P* < 0.0001.

**Figure S6.**
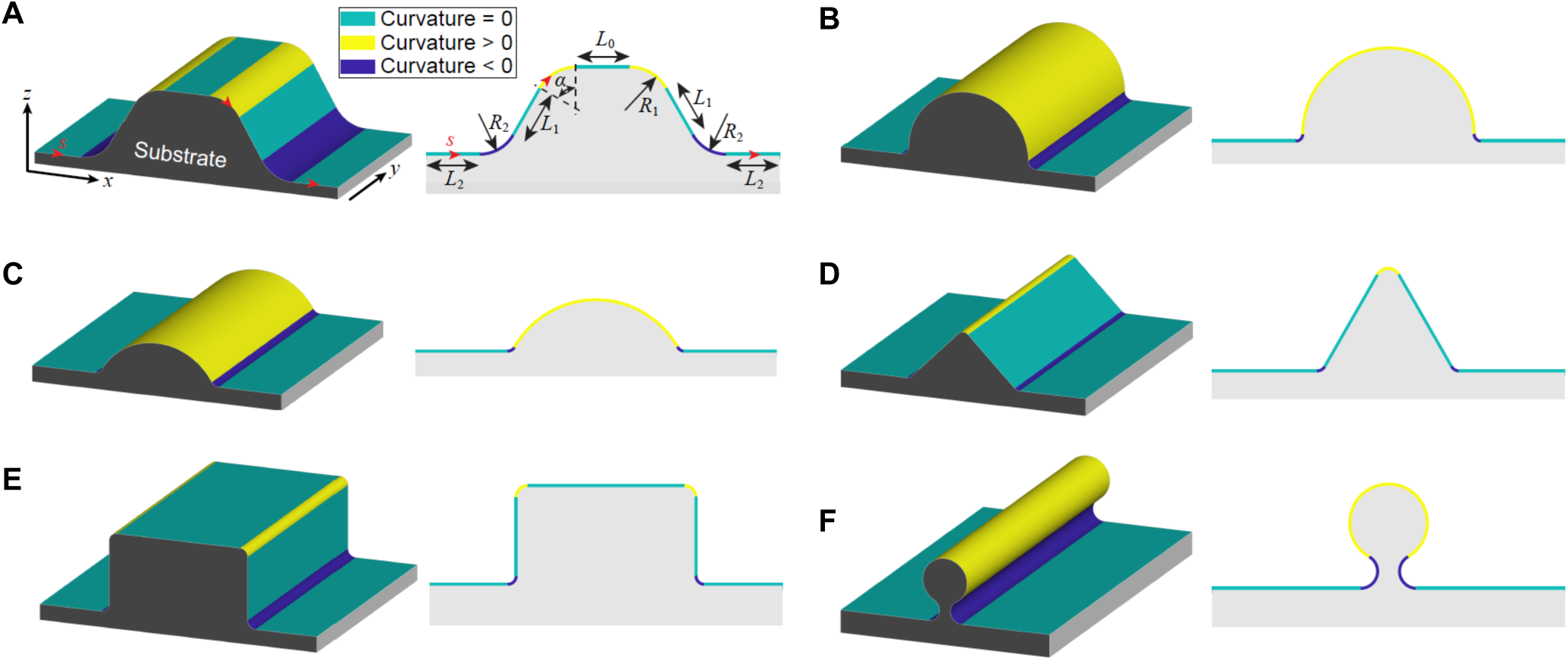
Illustration of *y*-invariant substrate geometries exhibiting modulated curvature. (A) Geometric description of the *y*-invariant curved substrate. It can be characterized by six parameters: *L*_0_, *L*_1_, *L*_2_, *R*_1_, *R*_2_, and *α.* In detail, the curved substrate is composed of nine regions, including two arcs of radius *R*_1_ and angle *α*, two arcs of radius *R*_2_ and angle *α*, one flat plane of length *L*_0_, two tilt planes of length *L*_1_, and two flat planes of length *L*_2_. (left) 3D view. Coordinate system: *y* in the invariant direction; curvilinear coordinate *s* in the plane normal to *y*; x in the horizontal direction, *z* in the vertical direction. (right) Side view. (B-F) Types of geometries mimicking our experiments. (B) Semi-cylinder with *L*_0_ = 0, *L*_1_ = 0, and *α* = π/2. (C) Cylindrical segment with smooth concave edges with *L*_0_ = 0, *L*_1_ = 0, and *α* < π/2. (D) Triangular prism with *L*_0_ = 0 and *α* < π/2. (E) Rectangular prism with *α* = π/2. (F) Nanofiber with *L*_0_ = 0, *L*_1_ = 0, and π/2 < *α* < π.

**Figure S7.**
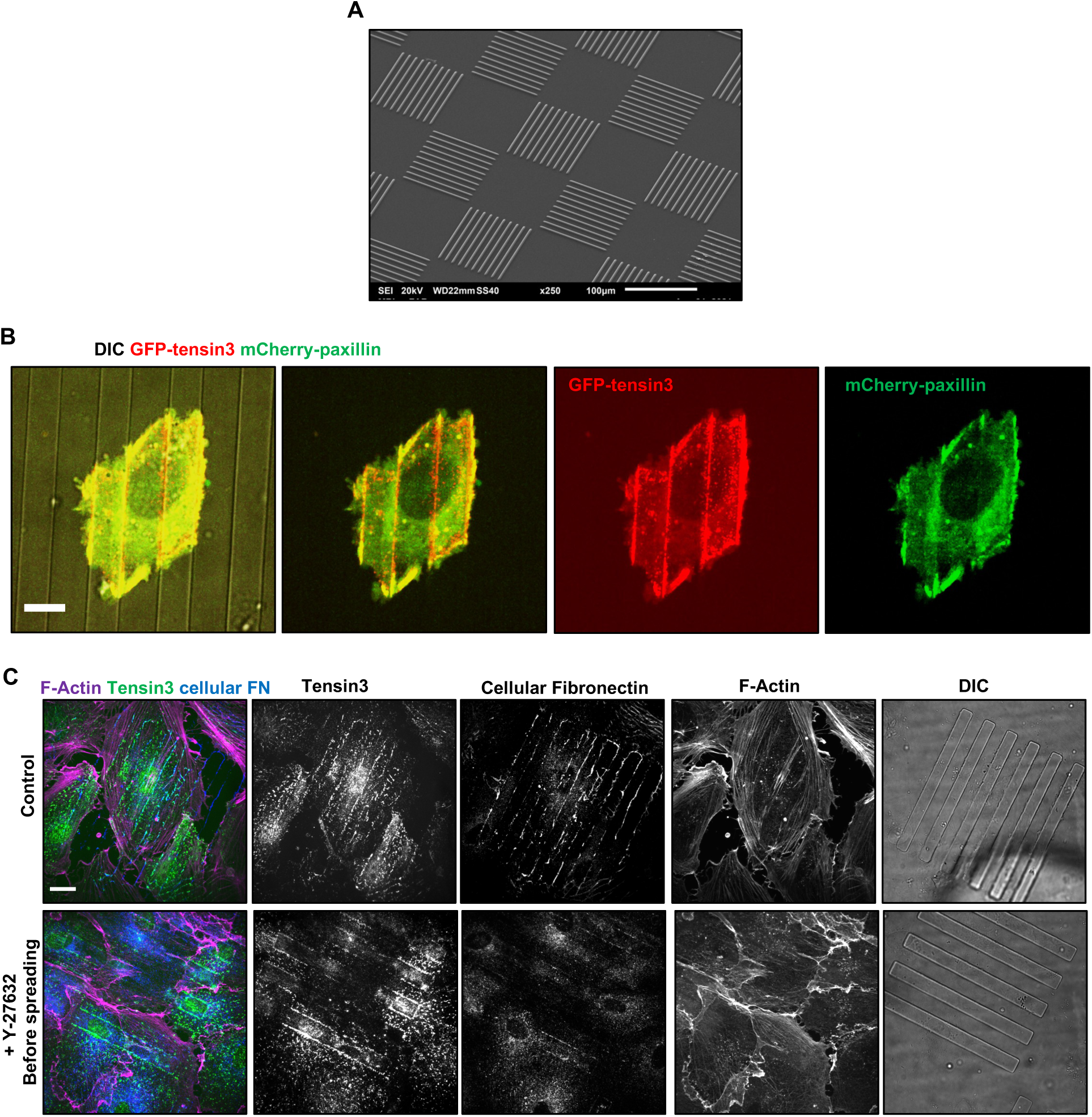
Localization of focal and fibrillar adhesions in cells grown on micro-ridges. (A) Scanning electron micrograph of a PDMS substrate containing semi-cylindrical micro-ridges with concave edges arranged in a chequerboard pattern. Scale bar, 100 µm. (B) Merged and single channel images of cells co-transfected with GFP–tensin3 (to label fibrillar adhesions, red) and mCherry–paxillin (to label focal adhesions, green) plated on plasma fibronectin-coated micro-ridges with a rectangular profile (visualized in DIC, grey). Tensin3-positive fibrillar adhesions form along concave micro-ridge edges while paxillin-positive focal adhesions form on the planar regions of the substrate. Scale bar, 10µm. (C) Images of cells grown on plasma fibronectin-coated micro-ridges with rectangular profile (DIC) in the absence or presence of 30 µM Y-27632. Left columns show merged images of F-actin (magenta), tensin3 (green), cellular fibronectin (blue) and right columns show individual channels including DIC to visualize the micro-ridges. Cells plated in the presence of ROCK inhibitor still form fibrillar adhesions along the edges of micro-ridges while fibronectin deposition and actin cytoskeleton organisation are perturbed. Scale bar, 10 µm.

**Figure S8.**
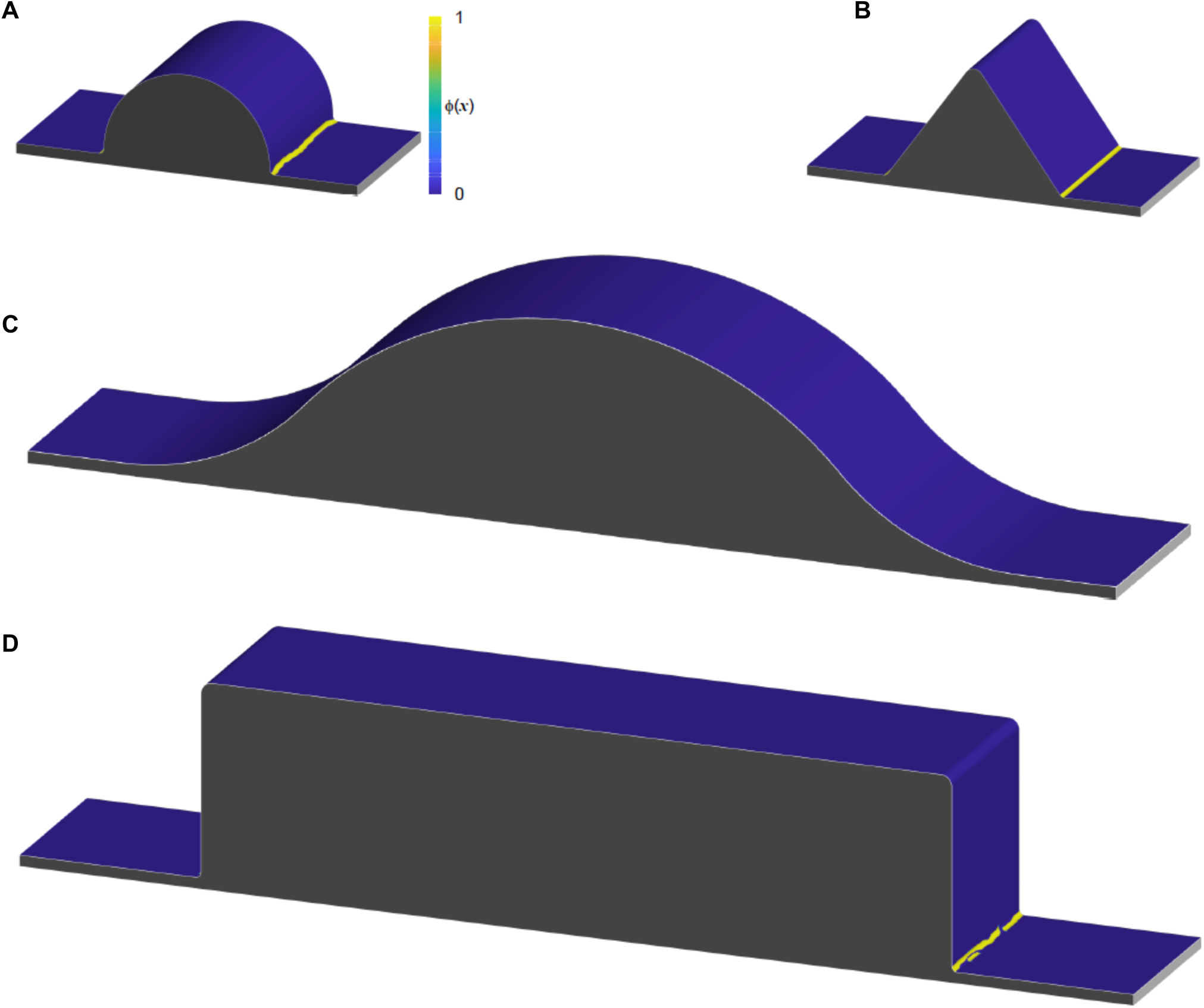
Numerical calculation of the clustering pattern of the fibrillar adhesion formation on curved substrates, driven by the anisotropic protein tilt effect. Numerical calculations over various curved substrates, including (A) the semi-cylinder geometry, (B) the triangle geometry, (C) the cylindrical segment geometry with smooth concave edges, and (D) the rectangle geometry. Parameters: *h_ϕ_* = 0.021 *k_B_T*·nm^−2^, σ = 0.076 *k_B_T*·nm^−2^, *ξ*_0_ = 0.0014 *k_B_T*·nm^−2^, and *ξ*_c_ = −0.0035 *k_B_T*·nm^−2^.

**Figure S9.**
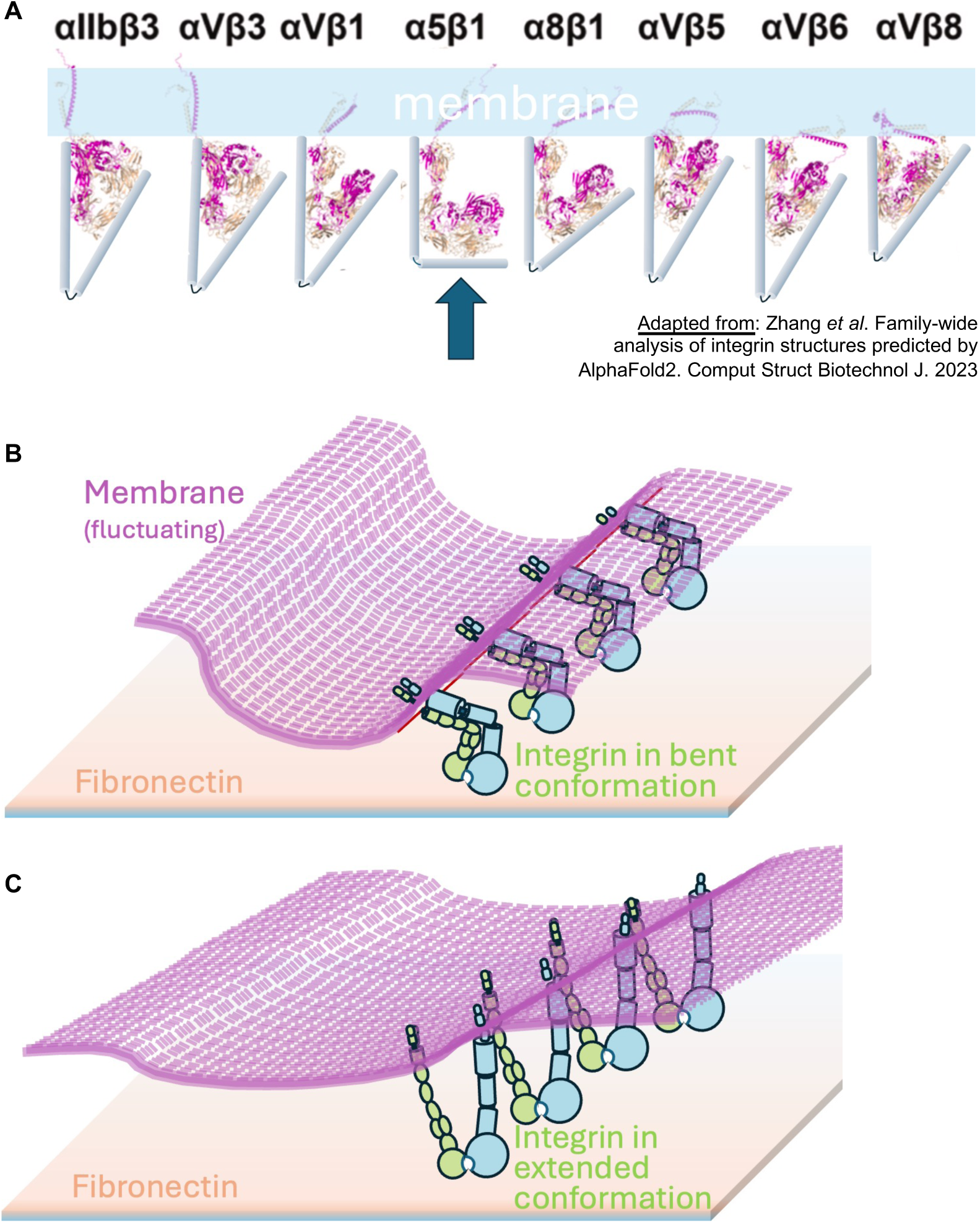
Special conformation of α5β1-integrin may facilitate its clustering along membrane folds and topographical cues. (A) AlphaFold2 reconstruction of the shape of extracellular domains of RGD-binding integrins in closed low affinity conformation (adapted from [57]). The alpha and beta subunits are shown in beige and magenta respectively. The integrins protrude from the membrane (blue) downwards. The transmembrane domain orientation was not correctly predicted for most of the structures [57]. The angles between two halves of the extracellular integrin moity are highlighted by blue “rods”. The shape of the extracellular domain of α5β1-integrin (arrow) is consistent with cryo-microscopy data [56] and differ from the predicted shapes of all other RGD binding integrins. While all the RGD-binding integrins have “V” configurations, in α5β1the two halves are approximately perpendicular to each other. (B and C) A hypothesis of α5β1-integrin clustering initiating the formation of fibrillar adhesions. (B) α5β1-integrin in bent conformation can undergo clustering along the membrane fold due to its unique shape (see also **Fig 7A**). (C) Upon binding to the ligand, the bent α5β1undergoes quick conformational changes (extension).

**Supplementary Movie 1**

Time-lapse movie of HUVEC plated on planar glass and co-transfected with mCherry–paxillin to which labels focal adhesions (magenta) and GFP–tensin3 which labels fibrillar adhesions (green). Merged images are on the left with single channel images of mCherry-paxillin in the middle and GFP-tensin3 on the right. Cells were treated with 30µM Y-27632 at 140s, resulting in the rapid shrinkage of paxillin-positive focal adhesions while no obvious effect on tensin3-positive central fibrillar adhesions is observed up to 12 min of imaging. Scale bar, 10µm.

**Supplementary Movie 2**

Time-lapse movie of FRAP experiment in HUVEC transfected with GFP–tensin3 and plated on electrospun nanofibers. The blue boxed regions mark nanofiber-associated GFP-tensin3 adhesions, and orange boxed regions mark adhesions on planar regions in between the nanofibers at: pre-bleach, bleach, and post-bleach recovery. Scale bar, 10µm.

**Supplementary Movie 3**

Time-lapse movie of HUVEC co-transfected with mCherry–myosin-IIA (magenta) and GFP–tensin3 (green) plated on electrospun nanofibers. Treatment with 1µM nocodazole (added just before imaging) for a total of 1 hour induced the disassembly of tensin3-positive fibrillar adhesions along nanofibers while GFP-tensin3-positive focal adhesions can be seeing growing along planar regions in between the nanofibers at the ends of myosin IIA filaments. Scale bar, 10µm.

**Supplementary Movie 4**

Time-lapse movie of HUVEC, plated on planar glass, transfected with GFP–tensin3 (green), localizing at centrally located, elongated, fibrillar adhesions as well as peripheral focal adhesion sites. Hypo-osmotic shock was induced at the start of imaging by diluting the culture medium 1:3 with distilled water (0.25X hypotonic). The movie shows disassembly of central fibrillar adhesions within 13 min of imaging with no apparent effect on peripheral focal adhesions. Scale bar, 10µm.

## Materials and Methods

### Cell culture

Human umbilical vein endothelial cells (HUVEC) (ATCC, CRL-1730) were cultured in endothelial growth medium EGM-2 Bulletkit (Lonza) including all provided supplements. Cells were kept at a passage less than 9, and incubated at 5% CO_2_ at 37°C. Primary human Foreskin Fibroblasts (HFF) (ATCC, SCRC-1041) were cultured in DMEM + 10% fetal bovine serum (FBS) and 1 mM sodium pyruvate, kept at an early passage (5-15) and incubated at 5% CO_2_ at 37°C.

### cDNA and RNAi transfection procedures

Plasmid transfection was carried out using electroporation (Neon Transfection System, Life Technologies) in accordance with the manufacturer’s instructions. Plasmids used were peGFP-IRSp53 (a kind gift from Dr Michael Sheetz lab), YFP-FCHo2 ([10] a kind gift from Dr. Wenting Zhao), peGFP-FCHo1 (addgene:196910), CIP4-mCherryC1 (addgene:27685), mCherry-paxillin [62], GFP-tensin1 and GFP-tensin3 (kind gifts from Dr. Benjamin Geiger), mCherry-myosin-IIA [63]. For siRNA transfection, cells were seeded at 75% confluency and transfected using RNAiMax (Life technologies) for siRNA (100 nM final concentration) following manufacturer’s instructions. Constructs used for RNAi were: ON-TARGETplus Non-targeting Control Pool, ON-TARGETplus Human MYH9 (4627) siRNA – SMARTpool, ON-TARGETplus Human MYH10 (4628) siRNA – SMARTpool, ON-TARGETplus Human FN1 (2335) siRNA – SMARTpool, ON-TARGETplus Human ITGA5 (3678) siRNA – SMARTpool, ON-TARGET plus Human TNS1 (7145) siRNA – SMARTpool, ON-TARGET plus Human TNS3 (64759) siRNA – SMARTpool.

### Drug treatments and hypo-osmotic shock

Pharmacological treatments were performed using the following concentrations of inhibitors or activators: 30 µM Y-27632 dihydrochloride (Sigma-Aldrich), 1 µg ml^−1^ Rho Activator II (CNO3, Cytoskeleton), 1 µM nocodazole (Sigma-Aldrich), 10mM Methyl-β-cyclodextrin (Sigma-Aldrich), 1 nM calyculin A (Sigma-Aldrich), 0.05M 1,6-Hexanediol (Sigma-Aldrich) or a similar concentration of DMSO (Sigma-Aldrich) as a vehicle control. The duration of the treatment with the inhibitors is indicated in figure legends. For hypotonic experiments, cells were exposed to solutions containing complete medium diluted in sterile distilled water at 1:3 (0.25× hypotonic) for 20 mins.

### Western Blotting

Protein lysates and immunoprecipitants were processed following standard procedures. Enhanced chemi-luminescence signal was acquired using an BioRad ChemiDoc Touch Imaging system. Antibody description and working concentrations can be found in **Table_S1**.

### Immunofluorescence

Cells were plated on 35mm glass bottom dishes (Iwaki) unless otherwise specified. Cells were fixed for 10 min with 4% paraformaldehyde (PFA) in PBS at room temperature, then permeabilized for 10 min using 0.2% Triton X-100 (Sigma-Aldrich) in PBS. Fixed cells were then blocked with 3% bovine serum albumin (BSA) for 1 hour at room temperature prior to incubation with primary antibodies **(Table_S1)** diluted in 3% BSA, overnight at 4°C. Samples were washed with 1XPBS three times and incubated with Alexa Fluor-conjugated secondary antibodies (Thermo Fisher Scientific) for 2 hours at room temperature, followed by three washes in 1XPBS. F-actin was visualized by Alexa Fluor 488 phalloidin (Thermo Fisher Scientific) or Alexa Fluor 647 phalloidin (Thermo Fisher Scientific).

**Table_S1.**

| Cat. No. | Antibody | Target | Manufacturer | Working concentrations |
| --- | --- | --- | --- | --- |
| #MABT201 | Anti-Integrin alpha5, clone SNAKA51 | α5-Integrin | Sigma Aldrich | I.F. (1:200) |
| ab150361 | Anti-Integrin alpha5 [EPR7854] | α5-Integrin | Abcam | W.B. (1:2000) |
| T5168 | Anti-TUBA4A (TUBA1) | αTubulin | Sigma Aldrich | W.B. (1:3000) |
| sc-376156 | Anti-Integrin alpha V/CD51 (H-2) | αv-Integrin | Santa Cruz | I.F. (1:100) |
| 552828 | Anti-Integrin beta1, clone mAB13 | β1-Integrin | BD Pharmingen | I.F. (1:100) |
| ab15459 | Anti-Integrin beta5 | β5-Integrin | Abcam | I.F. (1:100) |
| #610077 | Anti-Fibronectin, clone 10 | Fibronectin | BD Labs | I.F. (1:250), W.B. (1:2000) |
| ab2413 | Anti-Fibronectin | Fibronectin | Abcam | I.F. (1:250) |
| AB18265 | Anti-Fibronectin [Fn-3] | Fibronectin (cellular) | Abcam | I.F. (1:250) |
| sc-32233 | Anti-GAPDH (6C5) | GAPDH | Santa Cruz | W.B. (1:3000) |
| HPA015643 | Anti-KANK2 | KANK 2 | Atlas Antibodies | I.F. (1:100) |
| CMII 23 | Anti-Myosin IIB, non muscle | Myosin-IIB Heavy Chain | DSHB | W.B. (1:1000) |
| M8064 | Anti-Myosin IIA, non muscle | Myosin-IIA Heavy Chain | Sigma Aldrich | I.F. (1:250), W.B. (1:2000) |
| #610569 | Anti-Paxillin, clone 177 | Paxillin | BD Labs | I.F. (1:200) |
| ab241096 | Anti-Talin1 | Talin 1 | Abcam | I.F. (1:250) |
| HPA036089 | Anti-Tensin1 | Tensin 1 | Atlas Antibodies | I.F. (1:100), W.B. (1:2000) |
| #PA5-63112 | Anti-Tensin3 | Tensin 3 | Invitrogen | I.F. (1:100), W.B. (1:2000) |
| V9131 | Anti-Vinculin | Vinculin | Sigma Aldrich | I.F. (1:200) |
| ab71842 | Anti-Zyxin | Zyxin | Abcam | I.F. (1:200) |

### Confocal and Structured Illumination Microscopy (SIM)

After immunostaining, cells were imaged on a spinning-disc Nikon N-SIM microscope unless otherwise specified. This is based on a Nikon Ti-E inverted microscope with Perfect Focus System controlled by Nikon NIS-Elements AR software supplemented with a 100× oil immersion objective (1.40 NA, CFI Plan-ApochromatVC) and EMCCD camera (Andor Ixon DU-897).

### Single molecule localization microscopy

HUVECs were first plated on sterile coverslips containing immobilized gold nanoparticle fiducials. Cells were then fixed with 4% paraformaldehyde (Electron Microscopy Sciences) and 0.25% Triton X-100 (Sigma) in PHEM buffer for 15 min at 37 °C, followed by blocking in 3% BSA (Sigma) in PHEM buffer overnight at 4 °C. Samples were incubated with primary antibodies against tensin3 and α5-integrin (SNAKA51), followed by the respective secondary antibodies Alexa Fluor 488 (Invitrogen) and CF680 (Biotium). F-actin was visualized using Alexa Fluor 647–conjugated phalloidin (Invitrogen). Immediately before imaging, cells were mounted in an oxygen-scavenging imaging buffer containing 100 mM β-mercaptoethylamine (Sigma), 0.25 mg/mL glucose oxidase (Sigma), 20 µg/mL catalase (Sigma), and 2% (w/v) glucose in PHEM buffer. Alexa Fluor 488 fluorescence (tensin3) was used to locate regions of interest and was also recorded to allow overlap with single-molecule localizations of F-actin and α5-integrin.

Single molecule image acquisition was performed on an Abbelight SAFe 360 system built around an Olympus IX83 microscope equipped with two ORCA-FusionBT sCMOS cameras and a 100x oil immersion lens (UPlanApo, NA=1.5, Olympus). Spectral demixing was used to resolve Alexa Fluor 647 phalloidin and CF680-α5-integrin. During acquisition, the 647 laser was directed to uniformly excite samples in HILO mode while fluorescent emission of these two dyes was split by a long pass dichroic of 700 nm and then recorded simultaneously on two cameras. The histograms of intensity ratio [I1/(I1+I2)] of fluorescence signals at two cameras (I1, I2) from Alexa Fluor 647 and CF680 had narrow normal distributions with medians of 0.37 and 0.6, respectively, which showed negligible crosstalk and enabled simultaneous 2D dual-channel imaging of F-Actin (phalloidin) and α5-integrin (SNAKA51). A typical raw data set was composed of 20k sequential frames with an exposure time of 50ms for each camera. Raw data was further processed by custom-developed python software using 2D Gaussian function to localize the single molecule coordinates and reconstruct the super-resolved image[64]. Fiducials embedded on coverslips were used to register two channels at sub-pixel resolution and correct sample drift during acquisition.

### Fluorescence recovery after photobleaching (FRAP)

HUVEC were first transfected with GFP-tensin3 and plated on human plasma fibronectin-coated electrospun nanofibers cultured in EGM-2 medium at 37°C. FRAP experiments were performed on cells 24 hours post-transfection. The samples were mounted on a Nikon Eclipse Ti-E inverted microscope and viewed through a 100X objective (CFI Plan Apochromat λ 100X/1.45 NA oil immersion objective) coupled with a Yokogawa CSU-W1 Spinning Disk Field Scanning Confocal system. The entire setup was enclosed in a temperature-controlled environmental chamber maintained at 37°C. FRAP was executed using a computer-controlled iLas2 targeted laser illuminator, which selectively bleaches specified regions of interest (ROIs). For each cell, up to 12 ROIs were drawn surrounding individual fibrillar adhesions formed on either nanofibers or on the flat coverslip surface, each measuring 0.5 – 2.5 µm². Following the acquisition of several pre-bleach images, all ROIs within the field of view were simultaneously photobleached via a 150 mW pulse (∼9 s) using a 488 nm laser with ten repetitions for each ROI. Subsequent image acquisition was done at 250 ms intervals, each with an exposure time of 150 ms for a total duration of 1 minute post-photobleaching. Photoablation and image acquisition were controlled via MetaMorph software. Mean fluorescent intensity over time within each ROI were then quantified and normalized. The mobile fraction and τ values from each ROI were extracted using nonlinear regression analysis following an equation which either includes a single exponential term. Image sequences and data analysis were completed using Fiji and data visualizations were conducted through Python.

### Cell cortical stiffness measurement by atomic force microscopy (AFM)

The cortical stiffness [65, 66] of HUVEC were measured using a Nanowizard IV BioAFM system (JPK Instruments, Germany) on HUVEC cultured on plastic dishes and measured at 37 °C in EGM2 medium. Indentations were performed on randomly selected cells at the perinuclear region with a sharp tip (MLCT-C, Bruker, USA, nominal tip radius ∼20 nm). Force–distance curves were acquired at 5 μm/s with maximum force of 200 pN resulting in ∼200-400 nm indentations, within the thickness of the HUVEC cortex (∼400nm)[66]. Young’s modulus values were calculated using JPK Data Processing Software 6.4.21 (JPK Instruments, Germany), which employs Sneddon model for quadratic pyramid indenters (side angle 17.5°; nominal spring constant 0.01 N/m; Poisson’s ratio 0.5) fitted to the initial 80 nm of the approach curves. A sharp tip (small tip radius) is used as it concentrates stress in a smaller area and is more sensitive to the stiffness of the cortex (compared to spherical tip), hence useful in cortex-specific measurements [66]. HUVEC cells (control) and cells treated with the indicated drugs (30 µM Y-27632; 1 µM nocodazole; 1 nM calyculin A) were tested. HUVEC were incubated in Y-27632 and nocodazole for 60 mins before testing. Calyculin A was added directly to the HUVEC and the same cells were tested before and after 10 – 15 minutes of incubation. Cells showing excessive blebbing were omitted. An example brightfield image below shows the cantilever positioned over the cell before and after calyculin A treatment. Twenty five force curves per cell were characterized and averaged to evaluate the Young’s modulus for each condition.

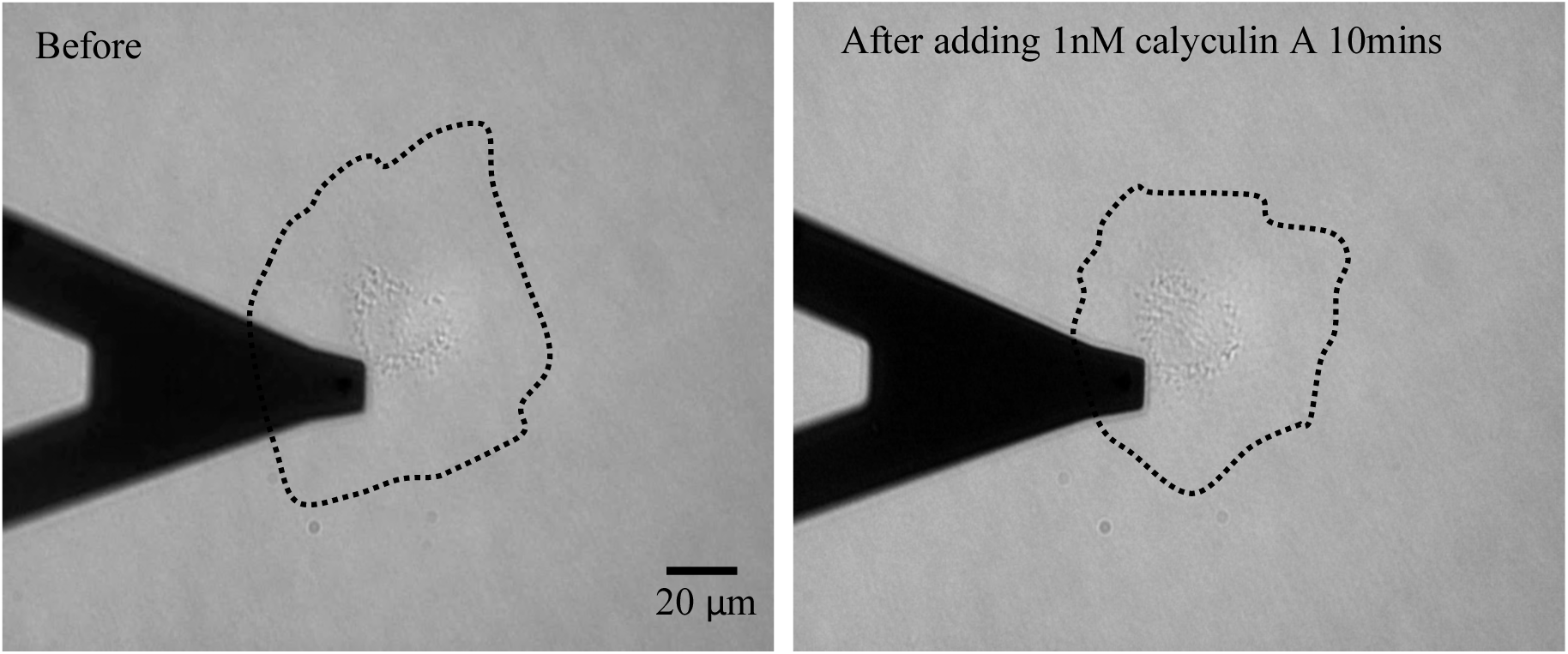

### Cell stretching assay

HUVEC were subjected to 16% rapid single stretch using the stretching device [67]. Briefly, cells were plated on a layer of polydimethylsiloxane (PDMS) coated with 10 µg ml^−1^ human plasma fibronectin (Merck), in a stretching unit. The substrate stretching was generated via changing the pressure in a chamber underneath the stretchable substrate. Cells were incubated under stretched conditions for 10 min, and then fixed as described above.

### Fabrication of PDMS micro-ridges (A-F)

Coverslips with PDMS micro-ridges were prepared by adapting a molding protocol from [68]. For each of the four geometries, a primary mold was microfabricated and coated with an anti-sticking layer. PDMS replicas were used as working molds to produce the coverslips.

#### (A) Silanization: anti-sticking coating with Trichloro(1H,1H,2H,2H-perfluorooctyl)silane (PFOCTS)

Trichloro(1H,1H,2H,2H-perfluorooctyl)silane (PFOCTS) (Sigma Aldrich CAS 78560-45-9) was used to prevent adhesion to the mold during replica casting, according to [69]. First, oxygen plasma activation was used to prepare the surface of the molds (20 sccm O_2_ flow at 2 mbar pressure in a Diener Pico tool, with either 60 W power at the RF generator for silicon moulds or 30 W power for PDMS mold, 1 min of process time); immediately after the plasma activation, the mold(s) were placed in a vacuum jar next to a small drop of PFOCTS (about 50 µL) and brought down to 1-2 mbar; after not less than 2 hr exposure to the vapours of PFOCTS, the mold(s) were retrieved ready for replica casting.

#### (B) PDMS casting – Replica molding

Sylgard 184 PDMS (Dow Corning) was prepared by mixing the elastomer base with its curing agent at a 10:1 ratio, thoroughly outgassed to remove any trapped air and poured on top of a silanized primary mold. To prevent defects possibly created by trapped air, a second outgassing in a vacuum jar (1-2 mbar for up to 30 min) was applied. The PDMS was then cured in an oven at 70 °C for 2 hr, diced in to final size and shape (e.g. 15 x 15 mm^2^ dices) and peeled-off from the primary mold.

#### (C) Primary mold for rectangular ridges

Rectangular cross-section features were fabricated using a one-step photolithography process. An optical mask with 8 µm width and 8 µm gap lines, 100 µm long arranged in patches of 100 × 100 µm^2^ was fabricated in house using a laser writer (Heidelberg DWL 66fs equipped with a 120 mW, 375 nm UV laser). A standard 4” silicon wafer was coated with a 2 µm thick layer of negative tone photoresist by spin-coating SU-8 2002 (Kayaku Advanced Materials, MA, USA) at 3000 rpm for 45” followed by soft baking on a hot plate (1 min at 65 °C followed by 1 min at 95 °C). The resist was exposed through the mask with UV light on a MJB4 mask aligner (SUSS MicroTec SE, Garching, DE), with a total energy dose of 80 mJ/cm2 at 354 nm wavelength. After post exposure baking (1 min at 65 °C followed by 1 min at 95 °C), the pattern was developed for 1 min in SU-8 developer (Kayaku Advanced Materials, MA, USA) rinsed with isopropyl alcohol (IPA), and dried with a nitrogen gun to reveal the line structures.

#### (D) Primary mold for semi-cylindrical ridges with concave edges and cylindrical segments with smooth concave edges

Ridges with semi-cylindrical or cylindrical cross-section were produced similarly. First, lines of 8 µm width and 2 µm height (for cylindrical segments) or 2 µm width and 1 µm height (for semi-cylindrical ridges) were produce by UV-lithography on a positive tone photoresist, AZ 5214E (AZ Electronic Materials). The photoresist was deposited by spin-coating on 4” silicon wafers, for 2 µm height at 1000 rpm for 45”; to produce the 1 µm thickness the photoresist was diluted in a 1:2 ratio by volume with his solvent (Propylene glycol methyl ether acetate, PGMEA, Kayaku Advanced Materials, MA, USA) prior to the spin-coating process. After pre-baking on a hot plate for 1 min at 120 °C the resist was exposed through a quartz optical mask for a total of 90 mJ/cm2 (at 354 nm wavelength) on a MJB4 mask aligner (SUSS MicroTec SE, Garching, DE). After 4 min development in AZ400K developer (AZ Electronic Materials) diluted 1:4 by volume in DI water, the wafers were placed back on the hot plate set at 140 °C and kept for 5 min to allow the photoresist to melt and reflow, which results in the curved profiles. After cooling down back to room temperature, the wafers were ready for the coating with PFOCTS.

#### (E) Primary mold for triangular cross sections

(100) SEMI2 standard silicon wafers with 300 nM of SiO_2_ thermally grown on both sides were spin-coated with approximately 1.4 µm thick AZ5214E positive tone photo-resist (spin coating at 2000 rpm followed by pre-baking on a hot plate at 110 °C for 1 min). Pattern of lines 2 µm wide were produced with direct writing in a DWL-66fs Heidelberg laser writer equipped with a diode-laser with 120 mW power at 375 nm. After development for 1 min in AZ400K diluted 1:4 in DI water, the patterned resist mask is then used to etch the silicon oxide layer in a Samco 10NR RIE tool using CF4/O2 etching chemistry (40/4 sccm respectively, 15 Pa, 150 W applied through an RF generator at 13.56 MHz, see [70] for more details). After the silicon oxide etching, the residual photo-resist mask was removed by oxygen plasma in the same reactor (40 sccm O_2_ flow, 10 Pa pressure, 90 W RF power, 10 min process time), then the wafer was immersed in 5M KOH at 80 °C, which produced trenches in the silicon with triangular section and by anisotropic wet etching. After anisotropic etching, the silicon oxide was removed with immersion in a buffered oxide etching (BOE 10:1, 901621-1L. Merck) for 10 min, followed with rinsing in DI water and drying with a nitrogen gun.

#### (F) Patterning of the glass coverslips

Glass coverslips were cleaned with soapy water, rinsed in DI water, then immersed in Acetone and rinsed in isopropyl alcohol and dried with a nitrogen gun. Prior to utilization, the cleaned glass coverslips were treated with oxygen plasma (20 SCCM O_2_ flow, 2 mbar pressure, 30 W Rf power and 1 min process time in a Harrick plasma tool), then immediately after spin-coated with PDMS (pre-mixed in 10:1 ratio and de-gassed) at 2000 rpm to produce a uniform 20-30 µm thick layer. A PDMS mold with the selected geometry for the ridges, silanized, was then placed on top of the coverslip and gently pressed. The coated coverslip with the mold were then placed in a vacuum chamber at about 1-2 mbar for 15-20 min, afterward PDMS was cured on a hot plate at 80°C for at least 1 hour. After curing, the PDMS mold was peeled-off revealing the patterned coverslip. Before cell plating, the PDMS substrates containing micro-ridges were coated with 10 µg/mL human plasma fibronectin (FC010 Merck) combined with 10 µg/mL Alexa Fluor 546 conjugated fibrinogen (Invitrogen, F13192) in PBS, for one-hour incubation at 37°C. During imaging, Z-stacks (0.1 µm step size) with the range set to encompass the crest and trough of the ridges were acquired. Three-dimensional reconstructions and orthogonal projections were done using Imaris.

### Preparation of thermoplastic polyurethane (TPU) nanofiber substrates

Twenty-five-millimeter round coverslips were cleaned in soapy water for 5 min, rinsed under running deionized (DI) water, and dried with a nitrogen gun. Subsequently, the coverslips were immersed in acetone (Thermo Fisher Scientific) for approximately 5 min, moved into Isopropyl alcohol (IPA) for another 5 min then rinsed with fresh IPA and dried using a nitrogen gun again. Right before the deposition of the nanofibers, clean coverslips were treated in oxygen plasma in a Tergeo Plasma Cleaner for 1 min at 20 W and 20 sccm O_2_ flow. The coverslips were then fixed to the rotating drum collector of an Inovenso NS1/Plus electrospinner. Nanofibers were generated from a polyurethane-based thermoplastic (TPU) solution (Inovenso) composed of 14% TPU, 70% dimethylformamide (DMF), and 16% ethyl acetate. Electrospinning was set for 15 s with 30 kV of applied voltage, 2.50 mL/hr TPU solution flow rate, with the drum collector rotating at 500 rpm and at a distance from the nozzle of 25 cm. Before cell plating, the nanofibers were coated with 10 µg/mL human plasma fibronectin (Merck), or 10 µg/mL laminin (Gibco), 20 µg/mL vitronectin (Invitrogen), 0.1% gelatin (Sigma) and incubated for 1 hr at 37°C. Coating solutions were then removed and the nanofiber substrates were washed once with 1× PBS before seeding cells at subconfluent density.

### Scanning electron microscopy (SEM) imaging

Scanning electron micrograph (SEM) images were acquired using a JEOL JSM-6010LV microscope. Non-conductive samples were prepared by coating with a thin (15 nm) metallic layer of Pt in a JEOL JFC-1600 Auto Fine coater, then loaded in the vacuum chamber of the SEM. Images were acquired at 20 keV acceleration of the electron gun (a W filament thermionic-emission gun).

### Image analysis

To quantify the relative area of fibrillar adhesions formed by cells on planar glass or on top of cell-derived matrix (CDM), images of fibrillar adhesions (typically α5-integrin or tensin) were thresholded in ImageJ/Fiji, and the total adhesion area was normalized to the total cell area (FB area / cell area). For semi-quantitative measurements of adhesion protein association along electrospun nanofibers, a nanofiber mask was generated from the corresponding DIC images. Fluorescent images of adhesion proteins were manually optimized to determine the mean adhesion intensity along nanofibers. The mask was then applied to extract the mean adjusted fluorescence intensity of the overlaid antibody staining, providing an estimate of adhesion occupancy on the nanofibers. For quantification of adhesion association with the underlying CDM, masks corresponding to the adhesion structures were first generated, and the mean fluorescence intensity for the underlying fibronectin (FN) staining within the CDM was measured. Co-localization analysis of myosin-IIA and SNAKA51 was performed using the JACoP plugin in ImageJ.

### Statistical analysis

Statistical analyses were performed using GraphPad Prism v10 (GraphPad Software, Inc.). The methods for statistical analysis and sizes of the samples (*n*) and individual biological replicates (*N*) are specified in the figure legends for all of the quantitative data. Differences were accepted as significant for *P* < 0.05. GraphPad version v10 or Python was used to plot and represent the data. Raw datasets generated during and/or analysed during the current study are available from the corresponding authors on reasonable request.

## Acknowledgments

We would like to thank Hui Ting Ong (NUS, Singapore) for help with image analysis, Giulia Adriani (A*STAR, Singapore) for providing HUVEC, Yee Han Tee (NUS, Singapore) for providing HFF, Mengya Kong (NTU, Singapore) for providing reagents and Corinne Albigez-Rizo (Université Grenoble Alpes, France) for helpful discussions. We also acknowledge the SIMBA Microscopy Facility and the Microfabrication Core Facility at the MBI for technical support. This work was supported by the Ministry of Education under the Research Centres of Excellence programme through the Mechanobiology Institute at the National University of Singapore (MBI core grant; ref. no. A-0003467-00-00) and by the National Research Foundation (NRF) Singapore, National University of Singapore, under its Mid-Sized Grant (NRF MSG; ref. no. NRF-MSG-2023-0001). A.D.B, and A.J.F are also supported by the Singapore Ministry of Education Academic Research Fund Tier 3 and Tier 2 grants (grant ref nos.: MOET32021-0003 and MOE-T2EP30223-0017). S.Z.L. is supported by the National Natural Science Foundation of China (Grant No. 12502367), Research Center for Magnetoelectric Physics of Guangdong Province (Grants 2024B0303390001), and Guangdong Provincial Key Laboratory of Magnetoelectric Physics and Devices (Grants 2022B1212010008).

## Author Contributions

A.J.F and A.D.B conceived the project and designed the experiments. A.J.F performed experiments and analysis with the assistance of D.Y.Y.J. and S.S.P. The theoretical model was designed and simulated by S.L, J.F.R and J.P, with input from A.J.F and A.D.B. STORM imaging and analysis was carried out by Y.W and P.K. AFM and analysis was carried out by N.M.H. H.C. carried out cell stretching experiment. S.F. and G.G. were responsible microfabrication of PDMS micro-ridges, electrospun nanofibers and SEM imaging. A.J.F and A.D.B prepared the manuscript with input from S.L, J.F.R and J.P.

## References

1. Bachmann, M., et al., Cell Adhesion by Integrins. Physiol Rev, 2019. 99(4): p. 1655–1699.

2. Winograd-Katz, S.E., et al., The integrin adhesome: from genes and proteins to human disease. Nat Rev Mol Cell Biol, 2014. 15(4): p. 273–88.

3. Fässler, R. and A. Sonnenberg, The integrin odyssey - a journey full of fundamental discoveries. J Cell Sci, 2025. 138(18).

4. Geiger, B., J.P. Spatz, and A.D. Bershadsky, Environmental sensing through focal adhesions. Nat Rev Mol Cell Biol, 2009. 10(1): p. 21–33.

5. Kanchanawong, P. and D.A. Calderwood, Organization, dynamics and mechanoregulation of integrin-mediated cell-ECM adhesions. Nat Rev Mol Cell Biol, 2023. 24(2): p. 142–161.

6. Zamir, E., et al., Molecular diversity of cell-matrix adhesions. J Cell Sci, 1999. 112 (Pt 11): p. 1655–69.

7. Linder, S., et al., Mechanisms and roles of podosomes and invadopodia. Nat Rev Mol Cell Biol, 2023. 24(2): p. 86–106.

8. Lock, J.G., et al., Reticular adhesions are a distinct class of cell-matrix adhesions that mediate attachment during mitosis. Nat Cell Biol, 2018. 20(11): p. 1290–1302.

9. Jacquemet, G., H. Hamidi, and J. Ivaska, *Filopodia in cell adhesion*, *3D* migration and cancer cell invasion. Curr Opin Cell Biol, 2015. 36: p. 23–31.

10. Zhang, W., et al., Curved adhesions mediate cell attachment to soft matrix fibres in three dimensions. Nat Cell Biol, 2023. 25(10): p. 1453–1464.

11. Cukierman, E., et al., Taking cell-matrix adhesions to the third dimension. Science, 2001. 294(5547): p. 1708–12.

12. Zamir, E., et al., Dynamics and segregation of cell-matrix adhesions in cultured fibroblasts. Nat Cell Biol, 2000. 2(4): p. 191–6.

13. Pankov, R., et al., Integrin dynamics and matrix assembly: tensin-dependent translocation of alpha(5)beta(1) integrins promotes early fibronectin fibrillogenesis. J Cell Biol, 2000. 148(5): p. 1075–90.

14. Fernandez-Sauze, S., et al., Regulation of fibronectin matrix assembly and capillary morphogenesis in endothelial cells by Rho family GTPases. Exp Cell Res, 2009. 315(12): p. 2092–104.

15. Atherton, P., et al., Tensin3 interaction with talin drives the formation of fibronectin-associated fibrillar adhesions. J Cell Biol, 2022. 221(10).

16. George, E.L., et al., Defects in mesoderm, neural tube and vascular development in mouse embryos lacking fibronectin. Development, 1993. 119(4): p. 1079–91.

17. Astrof, S. and R.O. Hynes, Fibronectins in vascular morphogenesis. Angiogenesis, 2009. 12(2): p. 165–75.

18. Huet-Calderwood, C., et al., Fibroblasts secrete fibronectin under lamellipodia in a microtubule- and myosin II-dependent fashion. J Cell Biol, 2023. 222(2).

19. Zhong, C., et al., Rho-mediated contractility exposes a cryptic site in fibronectin and induces fibronectin matrix assembly. J Cell Biol, 1998. 141(2): p. 539–51.

20. Lemmon, C.A., C.S. Chen, and L.H. Romer, Cell traction forces direct fibronectin matrix assembly. Biophys J, 2009. 96(2): p. 729–38.

21. Lu, J., et al., Basement Membrane Regulates Fibronectin Organization Using Sliding Focal Adhesions Driven by a Contractile Winch. Dev Cell, 2020. 52(5): p. 631–646.e4.

22. Baneyx, G., L. Baugh, and V. Vogel, Fibronectin extension and unfolding within cell matrix fibrils controlled by cytoskeletal tension. Proc Natl Acad Sci U S A, 2002. 99(8): p. 5139–43.

23. Clark, K., et al., A specific alpha5beta1-integrin conformation promotes directional integrin translocation and fibronectin matrix formation. J Cell Sci, 2005. 118(Pt 2): p. 291–300.

24. Barber-Perez, N., et al., Mechano-responsiveness of fibrillar adhesions on stiffness-gradient gels. J Cell Sci, 2020. 133(12).

25. Mould, A.P., S.K. Akiyama, and M.J. Humphries, The inhibitory anti-beta1 integrin monoclonal antibody 13 recognizes an epitope that is attenuated by ligand occupancy. Evidence for allosteric inhibition of integrin function. J Biol Chem, 1996. 271(34): p. 20365–74.

26. Heath, J.P. and G.A. Dunn, Cell to substratum contacts of chick fibroblasts and their relation to the microfilament system. A correlated interference-reflexion and high-voltage electron-microscope study. J Cell Sci, 1978. 29: p. 197–212.

27. Geiger, B., et al., Vinculin, an intracellular protein localized at specialized sites where microfilament bundles terminate at cell membranes. Proc Natl Acad Sci U S A, 1980. 77(7): p. 4127–31.

28. Friedl, K., et al., Assessing crosstalk in simultaneous multicolor single-molecule localization microscopy. Cell Reports Methods, 2023. 3(9): p. 100571.

29. Sottile, J. and D.C. Hocking, Fibronectin Polymerization Regulates the Composition and Stability of Extracellular Matrix Fibrils and Cell-Matrix Adhesions. Molecular Biology of the Cell, 2002. 13(10): p. 3546–3559.

30. Kaukonen, R., et al., Cell-derived matrices for studying cell proliferation and directional migration in a complex 3D microenvironment. Nat Protoc, 2017. 12(11): p. 2376–2390.

31. Elveren, B., et al., Cell Electrospinning: A Mini-Review of the Critical Processing Parameters and Its Use in Biomedical Applications. Advanced Biology, 2023. 7(10): p. 2300057.

32. Guo, K., et al., KANK1 shapes focal adhesions by orchestrating protein binding, mechanical force sensing, and phase separation. Cell Rep, 2023. 42(11): p. 113321.

33. Lee, Y.J., S. Yamada, and S.H. Lo, Phase transition of tensin-1 during the focal adhesion disassembly and cell division. Proc Natl Acad Sci U S A, 2023. 120(15): p. e2303037120.

34. Dibus, M., et al., Adhesion-derived condensates control component availability to regulate adhesion dynamics. bioRxiv, 2025: p. 2025.05.08.652869.

35. Li, X., et al., Talin–tensin3 interactions regulate fibrillar adhesion formation and tensin3 phase separation. Journal of Cell Biology, 2025. 225(1).

36. Alberti, S., A. Gladfelter, and T. Mittag, Considerations and Challenges in Studying Liquid-Liquid Phase Separation and Biomolecular Condensates. Cell, 2019. 176(3): p. 419–434.

37. Humphries, J.D., et al., *Molecular Basis of Ligand Recognition by Integrin α5β1: II. SPECIFICITY OF Arg-Gly-Asp BINDING IS DETERMINED BY Trp157* OF THE α SUBUNIT*. Journal of Biological Chemistry, 2000. 275(27): p. 20337–20345.

38. García, A.J., J.E. Schwarzbauer, and D. Boettiger, Distinct activation states of alpha5beta1 integrin show differential binding to RGD and synergy domains of fibronectin. Biochemistry, 2002. 41(29): p. 9063–9.

39. Rafiq, N.B.M., et al., Forces and constraints controlling podosome assembly and disassembly. Philos Trans R Soc Lond B Biol Sci, 2019. 374(1779): p. 20180228.

40. Truong Quang, B.A., et al., Extent of myosin penetration within the actin cortex regulates cell surface mechanics. Nat Commun, 2021. 12(1): p. 6511.

41. De Belly, H., et al., Cell protrusions and contractions generate long-range membrane tension propagation. Cell, 2023. 186(14): p. 3049–3061.e15.

42. Krendel, M., F.T. Zenke, and G.M. Bokoch, Nucleotide exchange factor GEF-H1 mediates cross-talk between microtubules and the actin cytoskeleton. Nature Cell Biology, 2002. 4(4): p. 294–301.

43. Biswas, A., et al., Cholesterol Depletion by MβCD Enhances Cell Membrane Tension and Its Variations-Reducing Integrity. Biophys J, 2019. 116(8): p. 1456–1468.

44. Cox, C.D., et al., Cyclodextrins increase membrane tension and are universal activators of mechanosensitive channels. Proc Natl Acad Sci U S A, 2021. 118(36).

45. Roffay, C., et al., Passive coupling of membrane tension and cell volume during active response of cells to osmosis. Proc Natl Acad Sci U S A, 2021. 118(47).

46. Lin, S.Z., et al., Membrane Tilt Drives Phase Separation of Adhesion Receptors. Phys Rev Lett, 2024. 132(18): p. 188402.

47. Litschel, T., et al., Membrane-induced 2D phase separation of the focal adhesion protein talin. Nat Commun, 2024. 15(1): p. 4986.

48. Kumar, A., K. Tanaka, and M.A. Schwartz, Focal adhesion-derived liquid-liquid phase separations regulate mRNA translation. Elife, 2025. 13.

49. Bershadsky, A.D., N.Q. Balaban, and B. Geiger, Adhesion-dependent cell mechanosensitivity. Annu Rev Cell Dev Biol, 2003. 19: p. 677–95.

50. Riveline, D., et al., Focal contacts as mechanosensors: externally applied local mechanical force induces growth of focal contacts by an mDia1-dependent and ROCK-independent mechanism. J Cell Biol, 2001. 153(6): p. 1175–86.

51. Ng, D.H., et al., Microtubule-dependent modulation of adhesion complex composition. PLoS One, 2014. 9(12): p. e115213.

52. Sitarska, E. and A. Diz-Muñoz, Pay attention to membrane tension: Mechanobiology of the cell surface. Curr Opin Cell Biol, 2020. 66: p. 11–18.

53. Kelkar, M., P. Bohec, and G. Charras, Mechanics of the cellular actin cortex: From signalling to shape change. Curr Opin Cell Biol, 2020. 66: p. 69–78.

54. Bruinsma, R. and E. Sackmann, Bioadhesion and the dewetting transition. Comptes Rendus de l’Académie des Sciences - Series IV - Physics-Astrophysics, 2001. 2(6): p. 803–815.

55. Sackmann, E. and R.F. Bruinsma, Cell adhesion as wetting transition? Chemphyschem, 2002. 3(3): p. 262–9.

56. Schumacher, S., et al., Structural insights into integrin α(5)β(1) opening by fibronectin ligand. Sci Adv, 2021. 7(19).

57. Zhang, H., D.S. Zhu, and J. Zhu, Family-wide analysis of integrin structures predicted by AlphaFold2. Computational and Structural Biotechnology Journal, 2023. 21: p. 4497–4507.

58. Li, J., J. Yan, and T.A. Springer, Low-affinity integrin states have faster ligand-binding kinetics than the high-affinity state. Elife, 2021. 10.

59. Li, J., et al., Ligand binding initiates single-molecule integrin conformational activation. Cell, 2024. 187(12): p. 2990–3005.e17.

60. Swaminathan, V., et al., Actin retrograde flow actively aligns and orients ligand-engaged integrins in focal adhesions. Proc Natl Acad Sci U S A, 2017. 114(40): p. 10648–10653.

61. Humphries, J.D., A. Byron, and M.J. Humphries, Integrin ligands at a glance. Journal of Cell Science, 2006. 119(19): p. 3901–3903.

62. Efimov, A., et al., Paxillin-dependent stimulation of microtubule catastrophes at focal adhesion sites. J Cell Sci, 2008. 121(Pt 2): p. 196–204.

63. Hu, S., et al., Long-range self-organization of cytoskeletal myosin II filament stacks. Nat Cell Biol, 2017. 19(2): p. 133–141.

64. Wang, W., et al., Chip-Based 3D Interferometric Nanoscopy. bioRxiv, 2025: p. 2025.05.13.653677.

65. Tee, S.Y., et al., Cell shape and substrate rigidity both regulate cell stiffness. Biophys J, 2011. 100(5): p. L25–7.

66. Vargas-Pinto, R., et al., The effect of the endothelial cell cortex on atomic force microscopy measurements. Biophys J, 2013. 105(2): p. 300–9.

67. Cui, Y., et al., Cyclic stretching of soft substrates induces spreading and growth. Nature Communications, 2015. 6(1): p. 6333.

68. Migliorini, E., et al., Acceleration of Neuronal Precursors Differentiation Induced by Substrate Nanotopography. Biotechnology and bioengineering, 2011. 108: p. 2736–46.

69. Schift, H., et al., Controlled co-evaporation of silanes for nanoimprint stamps. Nanotechnology, 2005. 16: p. 171–175.

70. Ashraf, M., S. Sundararajan, and G. Grenci, Low-power, low-pressure reactive-ion etching process for silicon etching with vertical and smooth walls for mechanobiology application. Journal of Micro/Nanolithography, MEMS, and MOEMS, 2017. 16(3): p. 034501.

