## Supplemental Theory for "Fibrillar adhesions are the primary integrin complexes shaped by matrix topography"

### Theory Notes

#### I. MODEL

To explain the clustering pattern of fibrillar adhesions on various curved substrates observed in our experiments, we consider a membrane-protein-substrate system. We describe such a system by two fields: (1) the distance between the membrane and the substrate, denoted $e(\boldsymbol{x})$; (2) the fraction of bound cell adhesion receptors, denoted $\phi(\boldsymbol{x}) \in (0, 1)$.

The stable clustering pattern of cell adhesion receptors is dictated by minimizing the free energy of the system, denoted $F$. We first provide the expression of $F$ for a generic curved substrate surface and then focus on a typical kind of curved surface, i.e., the $y$-invariant surface, which mimics the curved substrates used in our experiments. Following our previous work [1, 2], here, we consider mainly four contributions to the free energy of the system:

1. The entropy mixing energy $F_{\mathrm{FH}}$, which accounts for the mixing of cell adhesion receptors with binders. Following Refs. [1], we write the entropy mixing energy as:

$$F_{\mathrm{FH}} = \int \sqrt{g}\mathrm{d}\eta_1\mathrm{d}\eta_2 \left\{ \frac{k_BT}{a} \left[\phi \ln \phi + (1-\phi)\ln(1-\phi)\right] + \frac{1}{2}D_\phi (\nabla\phi)^2 \right\}, \tag{S1}$$

where $g = \det(g_{\alpha\beta})$ with $g_{\alpha\beta}$ the metric tensor of the membrane; $(\eta_1, \eta_2)$ is the surface coordinate defined on the substrate; $k_BT$ is the thermal energy; $a$ is the inverse areal density of binders, $D_\phi$ is a gradient energy coefficient; $\nabla$ represents the gradient differential operator within the membrane surface.

2. The mechanical energy of the membrane $F_{\mathrm{Hel}}$, which accounts for the elastic deformation of the membrane. We express $F_{\mathrm{Hel}}$ in the classical Helfrich formalism [3]:

$$F_{\mathrm{Hel}} = \int \sqrt{g}\mathrm{d}\eta_1\mathrm{d}\eta_2 \left[\sigma + \frac{1}{2}\kappa(c_\alpha^\alpha - c_0)^2\right], \tag{S2}$$

where $\sigma$ is the membrane tension; $\kappa$ is the bending stiffness of the membrane; $c_\alpha^\alpha$ is the mean curvature (defined as the sum of the two principal curvatures) of the membrane; $c_0$ is the spontaneous curvature of the membrane. We have shown that a coupling between $c_0$ and $\phi$ can lead to stable cluster formation [2]. Here, in our experiments, there is no evidence showing that such a spontaneous curvature mechanism is relevant for fibrillar adhesion formation. Thus, we set $c_0 = 0$ for simplicity.

3. The adhesion energy $F_{\mathrm{adh}}$, which accounts for the adhesion interactions between the membrane and the substrate. To the lowest order, we consider the following expression [1]:

$$F_{\mathrm{adh}} = \int \sqrt{g}\mathrm{d}\eta_1\mathrm{d}\eta_2 \left[\frac{1}{2}k_0e^2 - k_0e_0(1-\phi)e - h_\phi\phi\right], \tag{S3}$$

where $k_0 > 0$ is the membrane-substrate adhesion stiffness; $e_0$ stands for the rest-length distance between the membrane and the substrate; $h_\phi$ refers to the chemical potential of receptor-substrate binding.

4. The tilt energy $F_{\mathrm{tilt}}$, which accounts for the tilt effect of cell adhesion receptors with respect to the membrane or the substrate, as well as their bent conformation. For the fibrillar adhesions in our experiments, the cell adhesion receptors are integrins $\alpha_5\beta_1$, which have been reported to develop a tilt angle with respect to the membrane [4, 5]. Here, we consider a generic, anisotropic protein tilt effect. A generic free energy reads

$$F_{\mathrm{tilt}} = \int \sqrt{g}\mathrm{d}\eta_1\mathrm{d}\eta_2\phi \left\{\mu\theta_\alpha\nabla^{(s)\alpha}e + \frac{1}{2}\nu_{\alpha\beta}\theta^\alpha\theta^\beta\right\}, \tag{S4}$$

where $\overrightarrow{\theta} = \theta_\alpha\overrightarrow{\boldsymbol{e}}^{(s)\alpha}$ is the tilt vector describing the protein tilt direction and intensity with $\overrightarrow{\boldsymbol{e}}^{(s)\alpha}$ being a basis of contravariant vectors on the tangent plane of the substrate surface; $\nabla^{(s)}$ represents the gradient differential operator within the curved substrate surface; $\nu_{\alpha\beta}$ is an effective stiffness tensor that measures the cost of creating a tilt angle $\theta$. Minimizing the tilt energy Eq. (S4) leads to the following, simplified tilt energy:

$$F_{\mathrm{tilt}} = \int \sqrt{g}\mathrm{d}\eta_1\mathrm{d}\eta_2\phi \left\{-\frac{1}{2}\xi_{\alpha\beta}\nabla^{(s)\alpha}e\nabla^{(s)\beta}e\right\}, \tag{S5}$$

where the tensor $\xi_{\alpha\beta}$ quantifies the anisotropic protein tilt effect. The tensor $\xi_{\alpha\beta}$ is related to $\mu$ and $\nu_{\alpha\beta}$ as, $\xi_{\alpha\beta} = \mu^2(\nu^{-1})_{\alpha\beta}$ with $(\nu^{-1})^{\alpha\beta}$ being the inverse of $\nu_{\alpha\beta}$; see Sec. III B for details. We further take into account the fact that the substrate curvature can play the role of a field on the tilt orientation. To the lowest order, we consider a linear relation between the protein tilt $\xi_{\alpha\beta}$ and the substrate curvature $c^{(s)}_{\alpha\beta}$:

$$\xi_{\alpha\beta} = \xi_0 g^{(s)}_{\alpha\beta} + \xi_c c^{(s)}_{\alpha\beta}, \tag{S6}$$

where $\xi_0 > 0$ is the isotropic part and $\xi_c$ is the curvature-dependent part. When $\xi_c < 0$, the protein tilt effect is enhanced at concave substrate regions while suppressed at convex substrate regions. In contrast, when $\xi_c > 0$, the protein tilt effect is suppressed at concave substrate regions while enhanced at convex substrate regions. Note that, in the main text, we mainly focus on the case of isotropic tilt and discuss the anisotropic tilt effect in the discussion section. Our numerical calculations showed that the anisotropic tilt assumption with $\xi_c < 0$ predicts the exact clustering patterns of fibrillar adhesions for all curved substrate geometries used in our experiments (Fig. S8).

Therefore, the total free energy of the membrane-protein-substrate can be expressed as:

$$F = F_{\mathrm{FH}} + F_{\mathrm{Hel}} + F_{\mathrm{adh}} + F_{\mathrm{tilt}} = \int \sqrt{g}\mathrm{d}\eta_1 \mathrm{d}\eta_2 f[e,\phi], \tag{S7}$$

with

$$\begin{aligned} f[e,\phi] = {} & \frac{k_B T}{a}\left[\phi \ln \phi + (1-\phi)\ln(1-\phi)\right] + \frac{1}{2}D_\phi (\nabla\phi)^2 + \sigma + \frac{1}{2}\kappa(c^\alpha_\alpha - c_0)^2 \\ & + \frac{1}{2}k_0 e^2 - k_0 e_0 (1-\phi) e - h_\phi \phi - \frac{1}{2}\xi_{\alpha\beta}\phi \nabla^{(s)\alpha} e \nabla^{(s)\beta} e, \end{aligned} \tag{S8}$$

being the free energy density.

In particular, the substrate topographies in our experiments are invariant along a direction (here referred to as the $y$ direction), see Fig. S6 for examples.

Such $y$-invariant curved substrate surface is parameterized by $\eta_1 = s$ and $\eta_2 = y$ with $s$ being the arc length coordinate, see Fig. S6(a), with a curvature $c_s(s)$, i.e. the curvature of the intersection curve of the substrate surface and the plane $y = \mathrm{const}$. Consequently, the total free energy can be re-expressed as:

$$F = \int \mathrm{d}s\mathrm{d}y\sqrt{g} f(e;\phi), \tag{S9}$$

where

$$\begin{aligned} f(e;\phi) = {} & \frac{k_B T}{a}\left[\phi \ln \phi + (1-\phi)\ln(1-\phi)\right] + \frac{1}{2}D_\phi \left[g^{11}\left(\frac{\partial\phi}{\partial s}\right)^2 + 2g^{12}\frac{\partial\phi}{\partial s}\frac{\partial\phi}{\partial y} + g^{22}\left(\frac{\partial\phi}{\partial y}\right)^2\right] + \sigma + \frac{1}{2}\kappa(c^\alpha_\alpha - c_0)^2 \\ & + \frac{1}{2}k_0 e^2 - k_0 e_0 (1-\phi) e - h_\phi \phi - \frac{1}{2}\phi\left[(\xi_0 + \xi_c c_s)\left(\frac{\partial e}{\partial s}\right)^2 + \xi_0\left(\frac{\partial e}{\partial y}\right)^2\right], \end{aligned} \tag{S10}$$

with

$$g = g\left(e; \frac{\partial e}{\partial s}; \frac{\partial e}{\partial y}\right), \tag{S11}$$

$$g^{\alpha\beta} = g^{\alpha\beta}\left(e; \frac{\partial e}{\partial s}; \frac{\partial e}{\partial y}\right), \tag{S12}$$

$$c^\alpha_\alpha = c\left(e; \frac{\partial e}{\partial s}; \frac{\partial e}{\partial y}; \frac{\partial^2 e}{\partial s^2}; \frac{\partial^2 e}{\partial y^2}\right), \tag{S13}$$

as shown in Eqs. (S23)–(S25).

*Mathematical description of the y-invariant geometries.* – In general, we parameterize the curved substrate surface by: $\overrightarrow{OM}^{(s)}(\eta_1, \eta_2)$. With this parameterization, the covariant basis vectors of the substrate read:

$$\vec{\boldsymbol{e}}^{(s)}_{\alpha} = \frac{\partial \overrightarrow{OM}^{(s)}}{\partial \eta_\alpha}, \tag{S14}$$

which defines the normal vector of the substrate surface:

$$\vec{\boldsymbol{n}}^{(s)} = \frac{\vec{\boldsymbol{e}}^{(s)}_1 \times \vec{\boldsymbol{e}}^{(s)}_2}{|\vec{\boldsymbol{e}}^{(s)}_1 \times \vec{\boldsymbol{e}}^{(s)}_2|}. \tag{S15}$$

Then, we can describe the membrane position by:

$$\overrightarrow{OM}(\eta_1, \eta_2) = \overrightarrow{OM}^{(s)}(\eta_1, \eta_2) + e(\eta_1, \eta_2)\vec{\boldsymbol{n}}^{(s)}, \tag{S16}$$

where $e(\eta_1, \eta_2)$ is the distance between the membrane and the substrate. Further, using the same surface coordinates $(\eta_1,\, \eta_2)$ as the substrate, we can calculate the covariant basis vectors of the membrane by:

$$\vec{\boldsymbol{e}}_{\alpha} = \frac{\partial \overrightarrow{OM}}{\partial \eta_\alpha} = \vec{\boldsymbol{e}}^{(s)}_{\alpha} + \frac{\partial e}{\partial \eta_\alpha}\vec{\boldsymbol{n}}^{(s)} + e(\eta_1, \eta_2)\frac{\partial \vec{\boldsymbol{n}}^{(s)}}{\partial \eta_\alpha}, \tag{S17}$$

which defines the surface normal vector of the membrane as,

$$\vec{\boldsymbol{n}} = \frac{\vec{\boldsymbol{e}}_1 \times \vec{\boldsymbol{e}}_2}{|\vec{\boldsymbol{e}}_1 \times \vec{\boldsymbol{e}}_2|}. \tag{S18}$$

and the metric tensor of the membrane as,

$$g_{\alpha\beta} = \vec{\boldsymbol{e}}_{\alpha} \cdot \vec{\boldsymbol{e}}_{\beta}. \tag{S19}$$

The curvature tensor of the membrane reads:

$$\boldsymbol{c} = c^{\alpha\beta}\vec{\boldsymbol{e}}_{\alpha} \otimes \vec{\boldsymbol{e}}_{\beta} = c_{\alpha\beta}\vec{\boldsymbol{e}}^{\alpha} \otimes \vec{\boldsymbol{e}}^{\beta} \tag{S20}$$

where

$$c_{\alpha\beta} = \frac{\partial^2 \overrightarrow{OM}}{\partial \eta_\alpha \partial \eta_\beta} \cdot \vec{\boldsymbol{n}}, \tag{S21}$$

and $\{\vec{\boldsymbol{e}}^{\alpha}\}$ are the contravariant basis vectors, satisfying $\vec{\boldsymbol{e}}^{\alpha} \cdot \vec{\boldsymbol{e}}_{\beta} = \delta^{\alpha}_{\beta}$.

In particular, for the $y$-invariant curved substrate surface, see Fig. S6 for examples, we have:

$$(g_{\alpha\beta})_{2\times 2} = \begin{pmatrix} (1+c_s e)^2 + \left(\frac{\partial e}{\partial s}\right)^2 & \frac{\partial e}{\partial s}\frac{\partial e}{\partial y} \\ \frac{\partial e}{\partial s}\frac{\partial e}{\partial y} & 1 + \left(\frac{\partial e}{\partial y}\right)^2 \end{pmatrix} \tag{S22}$$

$$g = \det(g_{\alpha\beta}) = (1+c_s e)^2 \left[1 + \left(\frac{\partial e}{\partial y}\right)^2\right] + \left(\frac{\partial e}{\partial s}\right)^2 \tag{S23}$$

$$(g^{\alpha\beta})_{2\times 2} = \left[(g_{\alpha\beta})_{2\times 2}\right]^{-1} = \frac{1}{g}\begin{pmatrix} 1 + \left(\frac{\partial e}{\partial y}\right)^2 & -\frac{\partial e}{\partial s}\frac{\partial e}{\partial y} \\ -\frac{\partial e}{\partial s}\frac{\partial e}{\partial y} & (1+c_s e)^2 + \left(\frac{\partial e}{\partial s}\right)^2 \end{pmatrix} \tag{S24}$$

$$c^{\alpha}_{\alpha} = \frac{1}{g\sqrt{g}}\left\{\begin{array}{l} \left[1 + \left(\frac{\partial e}{\partial y}\right)^2\right]\left[-c_s(1+c_s e)^2 - 2c_s\left(\frac{\partial e}{\partial s}\right)^2 - e\frac{\partial e}{\partial s}\frac{\partial c_s}{\partial s} + (1+c_s e)\frac{\partial^2 e}{\partial s^2}\right] \\ + 2c_s\left(\frac{\partial e}{\partial s}\right)^2\left(\frac{\partial e}{\partial y}\right)^2 - 2\,(1+c_s e)\frac{\partial e}{\partial s}\frac{\partial e}{\partial y}\frac{\partial^2 e}{\partial s \partial y} \\ + \left[(1+c_s e)^2 + \left(\frac{\partial e}{\partial s}\right)^2\right](1+c_s e)\frac{\partial^2 e}{\partial y^2} \end{array}\right\}. \tag{S25}$$

#### II. NUMERICAL CALCULATION

##### A. Geometry

To mimic the curved substrates in our experiments, we consider a curved geometry shown in Fig. S6(a). Such a curved geometry is characterized by five geometric parameters: $L_0$, $L_1$, $L_2$, $R_1$, $R_2$, and $\alpha$. In detail, the curved substrate is composed of nine regions, including two arcs of radius $R_1$ and angle $\alpha$, two arcs of radius $R_2$ and angle $\alpha$, one flat plane of length $L_0$, two tilt planes of length $L_1$, and two flat planes of length $L_2$. By setting different parameters of ($L_0$, $L_1$, $L_2$, $R_1$, $R_2$, $\alpha$), we can mimic all the curved substrate geometries used in our experiments, including semi-cylinder, cylindrical segment, triangle, rectangle, and nanofibre, see Fig. S6(b-f). We list the geometric parameters in Table I and II.

##### B. Numerical scheme

To converge to the energy minimum, we consider the gradient-descent dynamics with an annealing procedure where the noise amplitude decays gradually with time [1, 2], i.e. we consider the evolution:

$$\partial e/\partial t = -\delta F/\delta e + \zeta(t), \tag{S26}$$

$$\partial \phi/\partial t = -\delta F/\delta \phi, \tag{S27}$$

where $\zeta(t)$ is Gaussian white noise. We perform numerical calculations in the $(s, y)$ space within a rectangular domain of size $L_s \times L_y$, using periodic boundary conditions. We use the spectral method to solve the controlling equations. The time integration is performed using a backward Euler scheme; the spatial derivatives are carried out using a second-order central difference method.

##### C. Parameter values

###### 1. *Geometric parameters*

According to our experiments, we set the geometric parameters ($L_0$, $L_1$, $L_2$, $R_1$, $R_2$, $\alpha$) as in Table I.

TABLE I. List of default geometric parameter values

| Parameter | Semi-cylinder | Cylindrical segment | Triangle | Rectangle | Nanofibre |
|---|---|---|---|---|---|
| $L_0$ | 0 | 0 | 0 | 8 $\mu$m | 0 |
| $L_1$ | 0 | 0 | 1.7 $\mu$m | 2 $\mu$m | 0 |
| $L_2$ | 1 $\mu$m | 1 $\mu$m | 1 $\mu$m | 2 $\mu$m | 1 $\mu$m |
| $R_1$ | 1 $\mu$m | 3.6 $\mu$m | 0.1 $\mu$m | 0.1 $\mu$m | 0.1 $\mu$m |
| $R_2$ | 0.05 $\mu$m | 2.4 $\mu$m | 0.05 $\mu$m | 0.05 $\mu$m | 0.05 $\mu$m |
| $\alpha$ | 90° | 47.2° | 54.5° | 90° | 150° |

###### 2. *Physical parameters*

According to previous studies, we use the parameter values as described in Table II. In our calculations, we normalize the parameters by the length scale $e_0 = 120$ nm [6–8] and the energy scale $k_BT = 4 \times 10^{-21}$J. We set the non-dimensional parameters as below: $\tilde{k}_0 = k_0 e_0^4/(k_BT) = 1000$, $\tilde{\kappa} = \kappa/(k_BT) = 10$, $\tilde{h}_\phi = h_\phi e_0^2/(k_BT) = 300$, $\tilde{a} = a/e_0^2 = 1/64$, and $\tilde{D}_\phi = D_\phi/(k_BT) = 1$.

TABLE II. List of default physical parameter values

| Parameter | Description | Value |
|---|---|---|
| $e_0$ | Length scale (membrane rest-length height) | 120 nm [6, 8] |
| $k_B T$ | Energy scale | $4 \times 10^{-21}$ J |
| $k_0$ | Membrane-substrate adhesion stiffness | $5 \times 10^{-6}\ k_B T \cdot \mathrm{nm}^{-4}$ [8] |
| $\kappa$ | Membrane bending stiffness | 10 $k_B T$ [8–11] |
| $a$ | Inverse areal density of binders | 225 $\mathrm{nm}^2$ [11, 12] |
| $D_\phi$ | Gradient energy coefficient | 1 $k_B T$ |

#### III. DISCUSSION ON THE SUBSTRATE CURVATURE DEPENDENT PROTEIN TILT EFFECT

##### A. Curvature-dependent protein tilt stiffness tensor

To show the protein tilt effect more intuitively, here, we consider the generalized tilt free energy described in Eq. (S4). We further assume that the protein tilt stiffness tensor $\nu_{\alpha\beta}$ depends on the substrate curvature $c^{(s)}_{\alpha\beta}$, which accounts for the case that the substrate curvature plays the role of a field acting on the protein tilt orientation. We consider a linear relation between $\nu_{\alpha\beta}$ and $c^{(s)}_{\alpha\beta}$:

$$\nu_{\alpha\beta} = \nu_0 g^{(s)}_{\alpha\beta} + \nu_c c^{(s)}_{\alpha\beta}, \tag{S28}$$

where $\nu_0$ is the isotropic part and $\nu_c$ is the curvature-dependent part.

For the $y$-invariant geometries in our experiments, the generalized tilt free energy Eq. (S4) further reads,

$$F_{\text{tilt}} = \int \mathrm{d}s\mathrm{d}y\sqrt{g} f_{\text{tilt}}, \tag{S29}$$

where

$$f_{\text{tilt}} = \phi\left\{\mu\left(\theta_s\frac{\partial e}{\partial s} + \theta_y\frac{\partial e}{\partial y}\right) + \frac{1}{2}\left(\nu_0 + \nu_c c_s\right)\theta_s^2 + \frac{1}{2}\nu_0\theta_y^2\right\}. \tag{S30}$$

To better illustrate how the substrate curvature affects the protein tilt configuration, here, we assume a constant gradient of membrane-substrate distance $(\partial e/\partial s, \partial e/\partial y) = (A\cos\psi, A\sin\psi)$ with $A$ the constant magnitude. Minimizing the tilt free energy leads to an optimal protein vector:

$$\theta_s^{(\text{opt})} = \pm\frac{\mu A}{(\nu_0 + \nu_c c_s)} \quad , \quad \theta_y^{(\text{opt})} = 0, \tag{S31}$$

if $\nu_c c_s < 0$; otherwise,

$$\theta_s^{(\text{opt})} = 0 \quad , \quad \theta_y^{(\text{opt})} = \pm\frac{\mu A}{\nu_0}. \tag{S32}$$

It suggests that, in the concave regions ($c_s < 0$), when $\nu_c > 0$, cell adhesion receptors prefer to tilt along the $s$ direction; whereas when $\nu_c < 0$, cell adhesion receptors prefer to tilt along the $y$ direction, i.e., the concave microridges. Figure TN1 shows the tilt free energy landscape for different cases of $\nu_c$.

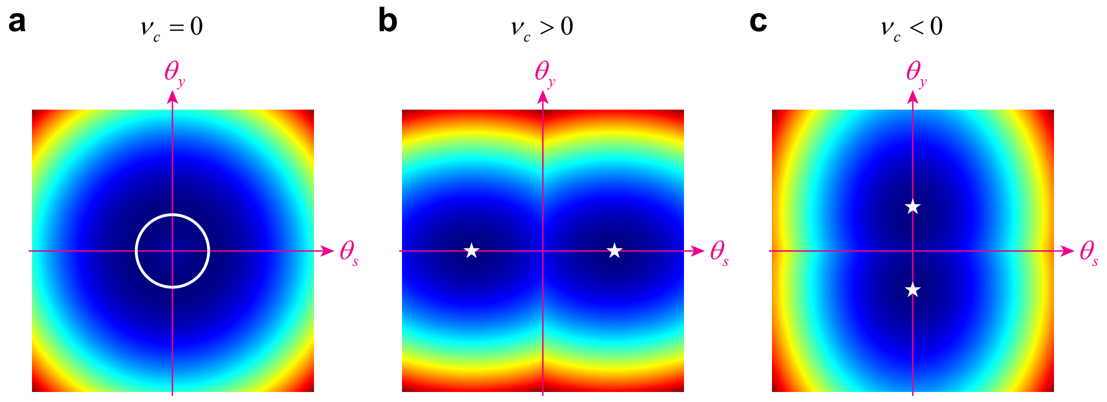


FIG. TN1. The tilt free energy Eq. (S30) as a function of the protein tilt vector $(\theta_s, \theta_y)$ in a concave region ($c_s < 0$), in the case of (a) $\nu_c = 0$, (b) $\nu_c > 0$, and (c) $\nu_c < 0$ (color bar: dark blue to red for increasing values). The white line or symbols indicate locations of the minima of the tilt free energy.

##### B. Relation to the curvature-dependent effective tilt tensor

For completeness, we now make explicit the connection between the generalized tilt model discussed in Sec. III A and the curvature-dependent effective tilt tensor $\xi_{\alpha\beta}$ introduced in Eq. (S6). Starting from the generalized tilt free energy, Eq. (S4), minimization with respect to the tilt vector $\theta^\alpha$ yields the optimal tilt configuration

$$\frac{\partial f_{\rm tilt}}{\partial \theta^\alpha} = 0 \quad \Rightarrow \quad \mu \nabla^{(s)}_\alpha e + \nu_{\alpha\beta}\theta^\beta = 0 \quad \Rightarrow \quad \theta^\alpha_{\rm opt} = -\mu(\nu^{-1})^{\alpha\beta}\nabla^{(s)}_\beta e, \tag{S33}$$

where $(\nu^{-1})^{\alpha\beta}$ denotes the inverse of the tensor $\nu_{\alpha\beta}$. Inserting $\theta^\alpha_{\rm opt}$ back into Eq. (S4) leads to

$$F_{\rm tilt} = \int \sqrt{g}\, d\eta_1 d\eta_2\, \phi \left\{ -\frac{1}{2}\mu^2(\nu^{-1})_{\alpha\beta}\nabla^{(s)\alpha} e \nabla^{(s)\beta} e \right\}, \tag{S34}$$

so that, by comparison with Eq. (S5), one can identify the effective tilt tensor as

$$\xi_{\alpha\beta} = \mu^2(\nu^{-1})_{\alpha\beta}. \tag{S35}$$

In Sec. III A, we have assumed that the protein tilt stiffness tensor $\nu_{\alpha\beta}$ itself depends on the substrate curvature; see Eq. (S28). For small curvature, one can expand the inverse tensor $(\nu^{-1})_{\alpha\beta}$ to linear order in $c^{(s)}_{\alpha\beta}$,

$$(\nu^{-1})_{\alpha\beta} \simeq \frac{1}{\nu_0} g^{(s)}_{\alpha\beta} - \frac{\nu_c}{\nu_0^2} c^{(s)}_{\alpha\beta} + \mathcal{O}\big((c^{(s)})^2\big). \tag{S36}$$

Combining this expansion with Eq. (S35) gives

$$\xi_{\alpha\beta} \simeq \mu^2 \left[ \frac{1}{\nu_0} g^{(s)}_{\alpha\beta} - \frac{\nu_c}{\nu_0^2} c^{(s)}_{\alpha\beta} \right]. \tag{S37}$$

Comparing with the phenomenological form introduced in Eq. (S6), one can identify, to lowest order in curvature,

$$\xi_0 \simeq \frac{\mu^2}{\nu_0}, \qquad \xi_c \simeq -\frac{\mu^2}{\nu_0^2}\nu_c. \tag{S38}$$

This relation shows that the curvature dependence of the effective tilt tensor $\xi_{\alpha\beta}$ used in Eq. (S6) is directly inherited from the curvature dependence of the microscopic stiffness tensor $\nu_{\alpha\beta}$. In particular, the sign of $\xi_c$ is opposite to the sign of $\nu_c$. As discussed in Sec. III A and illustrated in Fig. TN1, the sign of $\nu_c$ controls whether, in concave regions ($c_s < 0$), the receptors preferentially tilt along the $s$-direction or along the $y$-direction. Through the mapping above, this preference is encoded at the coarse-grained level by the sign of $\xi_c$ in Eq. (S6), which in turn determines how the substrate curvature locally modulates the effective tilt-mediated contribution to the membrane tension and hence the clustering of fibrillar adhesions.

---


[1] S.-Z. Lin, R. Changede, A. J. Farrugia, A. D. Bershadsky, M. P. Sheetz, J. Prost, and J.-F. Rupprecht, Physical Review Letters **132**, 188402 (2024).
[2] S.-Z. Lin, J. Prost, and J.-F. Rupprecht, Physical Review E **109**, 054406 (2024).
[3] W. Helfrich, Zeitschrift für Naturforschung C **28**, 693 (1973).
[4] J. Li, M. H. Jo, J. Yan, T. Hall, J. Lee, U. López-Sánchez, S. Yan, T. Ha, and T. A. Springer, Cell **187**, 2990 (2024).
[5] S. Schumacher, D. Dedden, R. V. Nunez, K. Matoba, J. Takagi, C. Biertümpfel, and N. Mizuno, Science Advances **7**, eabe9716 (2021).
[6] A.-S. Smith, K. Sengupta, S. Goennenwein, U. Seifert, and E. Sackmann, Proceedings of the National Academy of Sciences of the United States of America **105**, 6906 (2008).
[7] T. Bihr, U. Seifert, and A.-S. Smith, New Journal of Physics **17**, 083016 (2015).
[8] T. Bihr, U. Seifert, and A.-S. Smith, Physical Review Letters **109**, 258101 (2012).
[9] T. R. Weikl, Annual Review of Physical Chemistry **69**, 521 (2018).
[10] J. Steinkühler, E. Sezgin, I. Urbančič, C. Eggeling, and R. Dimova, Communications Biology **2**, 337 (2019).
[11] I. Raote, M. Chabanon, N. Walani, M. Arroyo, M. F. Garcia-Parajo, V. Malhotra, and F. Campelo, eLife **9**, e59426 (2020).
[12] X.-P. Xu, E. Kim, M. Swift, J. Smith, N. Volkmann, and D. Hanein, Biophysical Journal **110**, 798 (2016).
